# Tetraspanin Tspan15 is an essential subunit of an ADAM10 scissor complex

**DOI:** 10.1101/800557

**Authors:** Chek Ziu Koo, Neale Harrison, Peter J. Noy, Justyna Szyroka, Alexandra L. Matthews, Hung-En Hsia, Stephan A. Mueller, Johanna Tüshaus, Joelle Goulding, Katie Willis, Clara Apicella, Bethany Cragoe, Edward Davis, Murat Keles, Antonia Malinova, Thomas A. McFarlane, Philip R. Morrison, Michael C. Sykes, Haroon Ahmed, Alessandro Di Maio, Lisa Seipold, Paul Saftig, Eleanor Cull, Eric Rubinstein, Natalie S. Poulter, Stephen J. Briddon, Nicholas D. Holliday, Stefan F. Lichtenthaler, Michael G. Tomlinson

**Affiliations:** School of Biosciences, University of Birmingham, Birmingham, United Kingdom; Centre of Membrane Proteins and Receptors (COMPARE), Universities of Birmingham and Nottingham, Midlands, United Kingdom; German Center for Neurodegenerative Diseases (DZNE) Munich, Neuroproteomics, Klinikum rechts der Isar, Technical University Munich and Munich Cluster for Systems Neurology (SyNergy), Munich, Germany; Division of Physiology, Pharmacology and Neuroscience, School of Life Sciences, University of Nottingham, Nottingham, United Kingdom; Institute of Biochemistry, Christian Albrechts University Kiel, Kiel, Germany; Inserm, U935, F-94807, Villejuif, France; Institute of Cardiovascular Sciences, University of Birmingham, Birmingham, United Kingdom

**Author notes:** Authors contributed equally to this work (Chek Ziu Koo, Neale Harrison, Peter J. Noy). To whom correspondence should be addressed: Michael G. Tomlinson: School of Biosciences, College of Life and Environmental Sciences, University of Birmingham, Birmingham, B15 2TT, United Kingdom.

## Abstract

A disintegrin and metalloprotease 10 (ADAM10) is essential for embryonic development and impacts on diseases such as cancer, Alzheimer’s and inflammatory diseases. ADAM10 is a ‘molecular scissor’ that proteolytically cleaves the extracellular region from over 100 substrates, including Notch, amyloid precursor protein, cadherins, growth factors and chemokines. ADAM10 was recently proposed to function as six distinct scissors with different substrates, depending on its association with one of six regulatory tetraspanins, termed TspanC8s. However, it remains unclear to what degree ADAM10 function is critically dependent on a TspanC8 partner. To address this, we generated the first monoclonal antibodies to Tspan15 as a model TspanC8. These were used to show that ADAM10 is the principal Tspan15-interacting protein, that Tspan15 expression requires ADAM10 in cell lines and primary cells, and that a synthetic ADAM10/Tspan15 fusion protein is a functional scissor. Together these findings suggest that ADAM10 exists as an intimate ADAM10/TspanC8 scissor complex.

## Introduction

A disintegrin and metalloproteinase 10 (ADAM10) is a ubiquitously-expressed transmembrane protein which acts as a ‘molecular scissor’ by cleaving the extracellular region from over 100 substrates, a process termed ectodomain shedding (Lichtenthaler *et al.* 2018). ADAM10 is essential for embryonic development by activating Notch proteins which determine cell fate. Other substrates include cadherin adhesion molecules, amyloid precursor protein, and transmembrane growth factors and chemokines. As such, ADAM10 is important for health and in diseases such as cancer, Alzheimer’s and inflammatory diseases (Wetzel *et al.* 2017).

The tetraspanins are a superfamily of 33 transmembrane proteins in mammals which interact with specific transmembrane partner proteins and regulate their intracellular trafficking, lateral mobility and clustering at the cell surface (Termini and Gillette 2017; van Deventer *et al.* 2017). The first crystal structure of a tetraspanin suggests the capacity for conformational change, regulated by cholesterol binding within a cavity formed by the four transmembrane helices (Zimmerman *et al.* 2016). This raises the possibility that tetraspanins regulate their partner proteins as molecular switches. We and others have identified six TspanC8 tetraspanins as regulators of ADAM10 intracellular trafficking and enzymatic maturation (Dornier *et al.* 2012; Haining *et al.* 2012; Prox *et al.* 2012). The TspanC8s comprise Tspan5, 10, 14, 15, 17 and 33, and are so-called because of eight cysteines within their large extracellular loop. Different cell types express distinct TspanC8 repertoires (Matthews *et al.* 2017; Matthews *et al.* 2018) and emerging evidence suggests that each TspanC8 may cause ADAM10 to cleave specific substrates (Dornier *et al.* 2012; Jouannet *et al.* 2016; Noy *et al.* 2016; Reyat *et al.* 2017; Saint-Pol *et al.* 2017; Seipold *et al.* 2018; Brummer *et al.* 2019). However, there is an urgent need to generate monoclonal antibodies (mAbs) to all TspanC8s, to define fully their expression profiles and functional mechanisms. Moreover, it remains to be determined whether ADAM10 functions as an intimate ADAM10/TspanC8 complex or whether TspanC8s are merely modulators of ADAM10 trafficking.

This study focuses on Tspan15 as a model TspanC8. Tspan15 is best characterised by its capacity to promote ADAM10 cleavage of N-cadherin in cell lines and *in vivo* (Prox *et al.* 2012; Jouannet *et al.* 2016; Noy *et al.* 2016; Seipold *et al.* 2018). Tspan15 is also upregulated, and is a marker of poor prognosis, in certain cancers (Zhang *et al.* 2018; Hiroshima *et al.* 2019; Sidahmed-Adrar *et al.* 2019) and promotes cancer progression in a mouse model (Zhang *et al.* 2018). The aims of this study were to generate the first Tspan15 mAbs and to test three hypotheses that would support the theory that Tspan15 and ADAM10 exist together as a scissor complex: first, that ADAM10 is the principal Tspan15-interacting protein; second, that Tspan15 expression requires ADAM10; and third, that covalently linking Tspan15 and ADAM10 together as a single fusion protein yields a functional scissor.

## Results

### Generation of Tspan15 mAbs

The majority of anti-tetraspanin mAbs have epitopes within the large extracellular loop (LEL). However, it has been traditionally difficult to make mAbs to many tetraspanins due to lack of efficacy of recombinant LELs as immunogens. Furthermore, use of tetraspanins expressed in whole cells as the immunogen is complicated by their relatively high sequence conservation between species, their relatively small size and possible masking of mAb epitopes by larger partner proteins (Rubinstein *et al.* 2013). We therefore hypothesised that expression of human Tspan15 in ADAM10-knockout mouse cells would ‘unmask’ Tspan15, allowing the generation of a mAb response in mice immunised with these cells. Thus, ADAM10-knockout mouse embryonic fibroblasts (MEFs) (Reiss *et al.* 2005) stably overexpressing FLAG-tagged human Tspan15 were generated by lentiviral transduction, and cell lysates were immunoblotted with a FLAG antibody to confirm expression (Figure 1A). Immunisation of mice with these cells, and subsequent hybridoma generation, was outsourced to Abpro Therapeutics. Four positive hybridomas were identified by screening of hybridoma tissue culture supernatants by flow cytometry of wild-type versus CRISPR/Cas9 Tspan15-knockout Jurkat T cells (Figure 1B). The mAbs were isotyped as IgG1κ for clones 1C12, 4A4 and 5D4, and IgG2Bκ for 5F4 (data not shown). To investigate whether the Tspan15 mAbs might cross-react with other members of the TspanC8 family, HEK-293T cells were transfected with FLAG-tagged Tspan5, Tspan10, Tspan14, Tspan15, Tspan17 or Tspan33, and whole cell lysates western blotted. Tspan15 was detected by each of the four Tspan15 mAbs, but none of the other TspanC8s were detected (Figure 1C). To determine whether the Tspan15 mAbs can co-immunoprecipitate ADAM10, human platelets were lysed in the relatively stringent 1% digitonin lysis buffer that was previously used to identify TspanC8/ADAM10 interactions (Dornier *et al.* 2012; Haining *et al.* 2012). Tspan15 immunoprecipitation followed by ADAM10 western blotting showed that each mAb effectively co-immunoprecipitated ADAM10 (Figure 1D). Therefore, these data validate the four new mAbs as specific mouse anti-human Tspan15 reagents capable of immunoprecipitating Tspan15/ADAM10 complexes.

**Figure 1.**
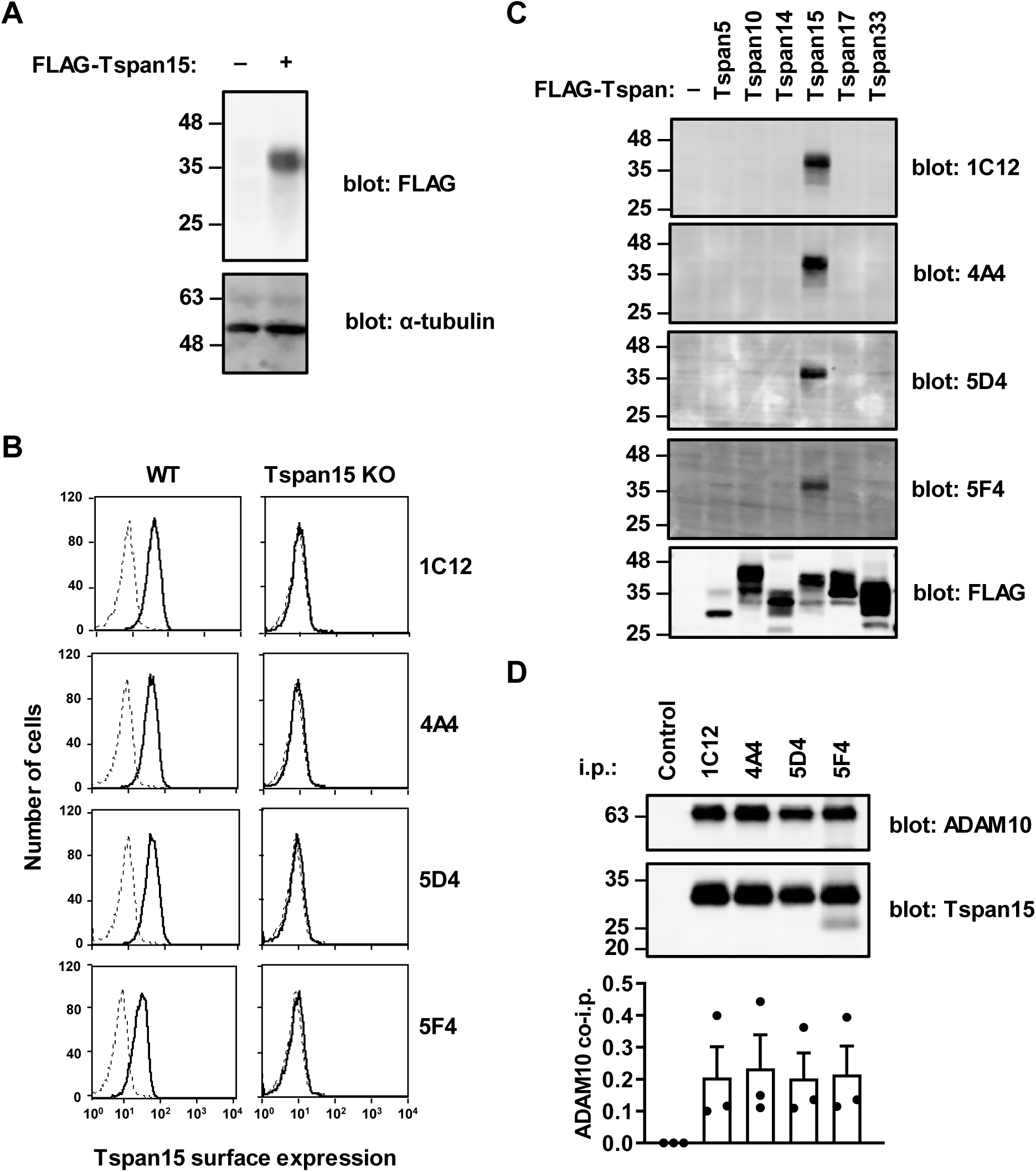
Generation of human Tspan15-expressing MEFs as an immunogen and validation of resulting mouse anti-human Tspan15 mAbs. (A) ADAM10-knockout MEFs (–) and ADAM10-knockout MEFs stably overexpressing FLAG-tagged Tspan15 (+) were lysed in 1% Triton X-100 lysis buffer and subjected to anti-FLAG (top panel) and anti-α-tubulin (bottom panel) western blotting. (B) Wild-type (WT) and Tspan15-knockout (KO) Jurkat human T cells were analysed by flow cytometry with tissue culture supernatant for each of the four mouse anti-human Tspan15 hybridomas (1C12, 4A4, 5D4 or 5F4; solid line), or with mouse IgG1 as a negative control (dotted line). Histograms are representative of two independent experiments. (C) HEK-293T cells were transfected with FLAG-tagged human TspanC8 expression constructs (except for Tspan10, which was of mouse origin) or an empty vector control (–), lysed in 1% Triton X-100 lysis buffer and western blotted with tissue culture supernatants for each of the four Tspan15 hybridomas, or a positive control FLAG antibody. Blots are representative of three independent experiments. (D) Human platelets were lysed in 1% digitonin lysis buffer and subjected to immunoprecipitation with the four Tspan15 mAbs or a negative control mouse IgG1, followed by anti-ADAM10 and anti-Tspan15 (5D4) western blotting (upper panels). The faint additional band in the 5F4 lane corresponds to light chain from the immunoprecipitating mAb (data not shown). To quantitate the data, the amount of ADAM10 co-immunoprecipitated was normalised to the amount of immunoprecipitated Tspan15 with each antibody (lower panel). Error bars represent standard error of the mean from three independent experiments.

### The four Tspan15 mAbs bind to similar epitopes in the large extracellular loop

To determine whether the Tspan15 mAbs have epitopes in the large extracellular loop (LEL), previously described Tspan15/Tspan5 chimeric GFP-tagged constructs were used in which the LELs were exchanged (Saint-Pol *et al.* 2017). In flow cytometry analyses of transfected HEK-293T cells, the four Tspan15 mAbs detected the Tspan15 LEL on a Tspan5 backbone (T5-LEL15), but not the reciprocal protein (T15-LEL5) (Figure 2A). As a control, a Tspan5 mAb detected Tspan5 LEL on a Tspan15 backbone (Figure 2A), as previously reported (Saint-Pol *et al.* 2017). To investigate whether the Tspan15 mAbs have overlapping epitopes, an antibody binding competition assay was performed. To do this, the A549 lung epithelial cell line was preincubated with one of the Tspan15 mAbs or a control antibody, and then incubated with Tspan15 mAbs conjugated to Alexa Fluor® 647. Flow cytometry analyses showed that all four unlabelled Tspan15 mAbs inhibited binding of the each of the four labelled Tspan15 mAbs (Figure 2B), suggesting that their epitopes are located in close proximity. Within the Tspan15 LEL, only eight of the 121 amino acid residues vary between mouse and human, all within the C-terminal half of the LEL (Figure 2C). Since none of the Tspan15 mAbs detected mouse Tspan15 by western blotting (Figure 2D), this suggested that some of the differing residues were important for antibody binding. To determine which were important, four FLAG-tagged Tspan15 human/mouse chimeric constructs were made by substituting residues within the LEL of human Tspan15 with the corresponding mouse residues (Figure 2C), an approach that we used previously to epitope map mAbs to tetraspanin CD53 (Tomlinson *et al.* 1993; Tomlinson *et al.* 1995). Western blotting of lysates from transfected Tspan15-knockout HEK-293T cells showed that reactivity of all four Tspan15 mAbs was almost completely lost by changing residues FSV to LNA, but unaffected in the three other chimeras (Figure 2D). However, the human FSV sequence was not sufficient for recognition, because introduction of these residues into mouse Tspan15 did not enable recognition by the four Tspan15 mAbs (Figure 2E). Taken together, these data show that all four Tspan15 mAbs bind to similar epitopes within the LEL.

**Figure 2.**
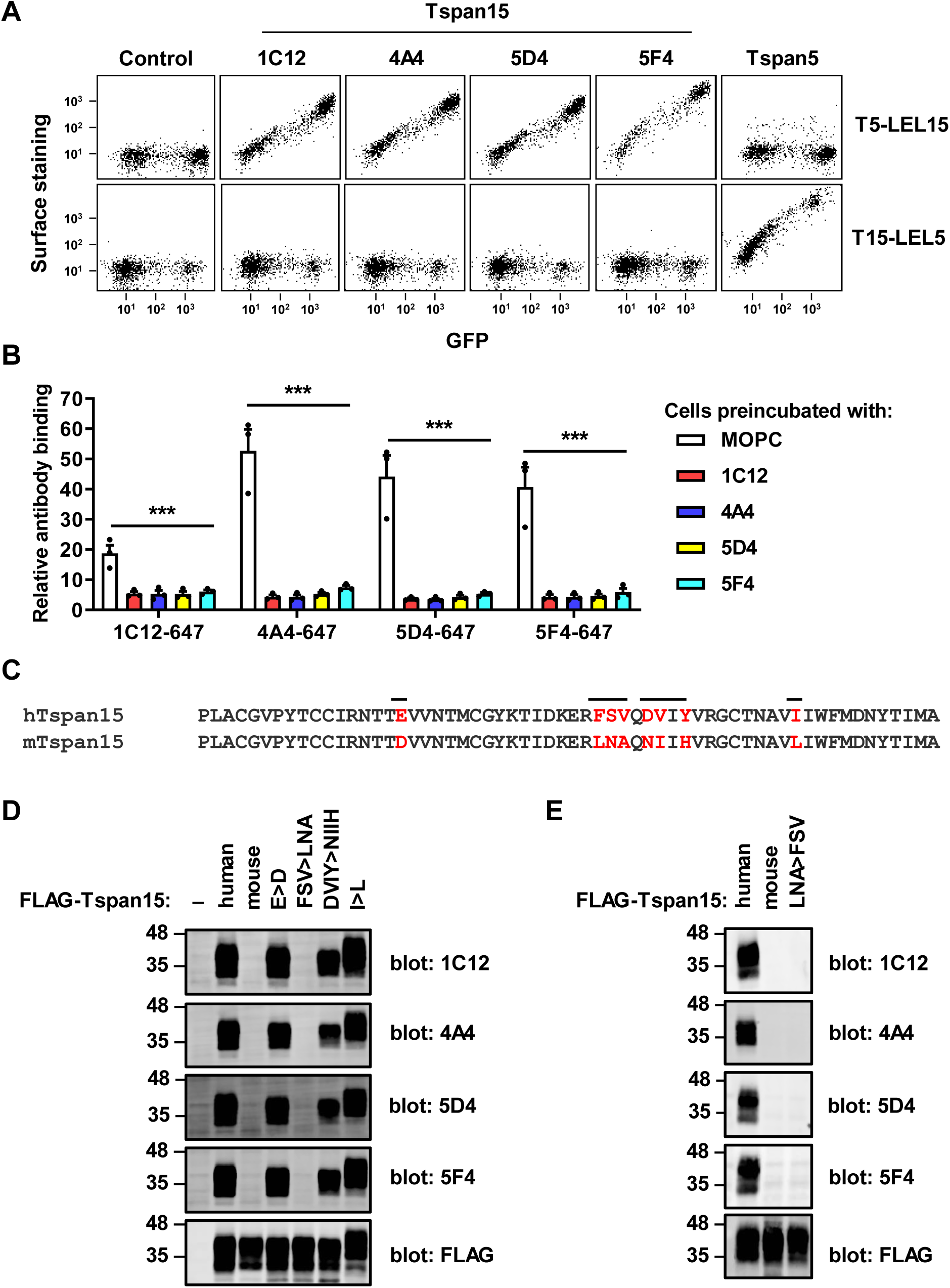
The four Tspan15 mAbs bind to similar epitopes in the large extracellular loop. (A) Tspan15-knockout HEK-293T cells were transfected with GFP-tagged Tspan5 with the large extracellular loop of Tspan15 (T5-LEL15) or the reciprocal chimeric expression construct (T15-LEL5). Cells were stained with the four Tspan15 mAbs (1C12, 4A4, 5D4 or 5F4), Tspan5 mAb (TS5-2) or negative control mouse IgG1, followed by APC-conjugated anti-mouse antibody and flow cytometry analyses. (B) A549 cells were preincubated with one of the four unlabelled Tspan15 mAbs, or a negative control mouse IgG1, for 30 minutes and stained with Alexa Fluor® 647-conjugated Tspan15 mAbs. Antibody binding, relative to unstained cells, was quantitated by flow cytometry. Data were log-transformed and statistically analysed by a two-way ANOVA with Tukey’s multiple comparisons test. Error bars represent the standard error of the mean from three independent experiments (\*\*\**p*<0.001 for control compared to each of the mAb preincubations). (C) Amino acid sequence alignment of the C-terminal half of human (h) and mouse (m) Tspan15 large extracellular loop region with Clustal Omega (Sievers *et al.* 2011). Sequence differences are in red, and sequences exchanged in the four mutant constructs are indicated by horizontal lines above the line-up. (D) Tspan15-knockout HEK-293T cells were transfected with FLAG-tagged human Tspan15, mouse Tspan15, four chimeric constructs with human Tspan15 residues replaced by corresponding mouse residues, or an empty vector control (−). Cells were lysed in 1% Triton X-100 lysis buffer and subjected to anti-Tspan15 or anti-FLAG western blotting. (E) Tspan15-knockout HEK-293T cells were transfected with FLAG-tagged human Tspan15, mouse Tspan15 or a chimeric construct comprising mouse Tspan15 with three residues of the corresponding human sequence. Cell lysates were western blotted as described in panel D. Data for panels D-E are each representative of three experiments, from which quantitation demonstrated no significant detection of mouse Tspan15 or the FSV to LNA and reciprocal chimera (data not shown).

### ADAM10 is the principal Tspan15-interacting protein

The generation of Tspan15 mAbs enabled the first proteomic identification of interactors with endogenous Tspan15. To do this, Tspan15 mAb 1C12 was used to immunoprecipitate Tspan15 from wild-type HEK-293T cells lysed in the relatively stringent detergent 1% digitonin; Tspan15-knockout cells were used as a negative control. Subsequent mass spectrometry identification revealed 28 proteins that were significantly detected in wild-type versus Tspan15-knockout samples from five independent experiments (Table 1 and Supplementary Table 1). Expression of the entire dataset as a volcano plot illustrated how the most significant and differential protein identified was ADAM10 (Figure 3). Indeed, ADAM10 and Tspan15 were the only proteins above the false discovery threshold for these experiments (Figure 3). These data suggest that ADAM10 is the principal Tspan15-interacting protein in HEK-293T cells.

**Table 1.** Proteins identified in Tspan15 immunoprecipitates. The table contains proteins significantly enriched in the Tspan15 immunoprecipitation samples of wild-type (WT) compared to Tspan15-knockout (KO) samples. Five additional proteins detected in WT samples (in at least 3 of 5 biological replicates), but not in more than one Tspan15 KO sample, are indicated with an asterisk. UniProt accession, protein name, gene name, *p*-value, fold difference between wild-type and Tspan15-KO samples are listed. In addition, UniProt annotations indicate the localization of the proteins.

| Accession | Protein name | Gene name | *p*-value | Fold difference (WT vs KO) | Localization |
| --- | --- | --- | --- | --- | --- |
| **O14672** | Disintegrin and metalloproteinase domain-containing protein 10 | ADAM10 | 1.70E-08 | 397.32 | Membrane |
| **O95858** | Tetraspanin-15 | TSPAN15 | 3.38E-05 | 18.57 | Membrane |
| **P52209** | 6-phosphogluconate dehydrogenase, decarboxylating | PGD | 2.28E-02 | 4.18 | Cytoplasm |
| **P62879** | Guanine nucleotide-binding protein G(I)/G(S)/G(T) subunit beta-2 | GNB2 | 2.80E-02 | 3.02 | Cytoplasm |
| **Q08380** | Galectin-3-binding protein | LGALS3BP | 2.92E-03 | 2.92 | x |
| **Q14315** | Filamin-C | FLNC | 1.10E-02 | 2.34 | Cytoplasm, Membrane |
| **P46779** | 60S ribosomal protein L28 | RPL28 | 3.47E-02 | 1.89 | x |
| **P27824** | Calnexin | CANX | 1.82E-02 | 1.86 | Membrane |
| **Q8WU90** | Zinc finger CCCH domain-containing protein 15 | ZC3H15 | 1.51E-02 | 1.75 | Cytoplasm |
| **Q8NEZ5** | F-box only protein 22 | FBXO22 | 1.02E-02 | 1.70 | Cytoplasm |
| **Q9HAD4** | WD repeat-containing protein 41 | WDR41 | 2.05E-02 | 1.61 | Cytoplasm |
| **Q12907** | Vesicular integral-membrane protein VIP36 | LMAN2 | 1.97E-02 | 1.59 | Membrane |
| **P63000** | Ras-related C3 botulinum toxin substrate 1 | RAC1 | 1.96E-02 | 1.53 | Cytoplasm, Membrane |
| **P23528** | Cofilin-1 | CFL1 | 3.95E-02 | 1.45 | Cytoplasm, Membrane |
| **Q4KMP7** | TBC1 domain family member 10B | TBC1D10B | 3.35E-03 | 1.40 | Cytoplasm |
| **Q9Y2G5** | GDP-fucose protein O-fucosyltransferase 2 | POFUT2 | 3.52E-02 | 1.35 | x |
| **P36639** | 7,8-dihydro-8-oxoguanine triphosphatase | NUDT1 | 2.69E-02 | 1.35 | Cytoplasm |
| **Q15020** | Squamous cell carcinoma antigen recognized by T-cells 3 | SART3 | 4.79E-02 | 1.34 | Cytoplasm |
| **Q9P2E9** | Ribosome-binding protein 1 | RRBP1 | 2.95E-02 | 1.33 | Membrane |
| **Q13042** | Cell division cycle protein 16 homolog | CDC16 | 2.39E-02 | 1.31 | Cytoplasm |
| **P27348** | 14-3-3 protein theta | YWHAQ | 2.66E-02 | 1.27 | Cytoplasm |
| **Q8WUM4** | Programmed cell death 6-interacting protein | PDCD6IP | 1.56E-02 | 1.24 | Cytoplasm |
| **P40926** | Malate dehydrogenase, mitochondrial | MDH2 | 3.49E-02 | 1.18 | x |
| **P62304** | Small nuclear ribonucleoprotein E | SNRPE | * | 0.73 | Cytoplasm |
| **O43172** | U4/U6 small nuclear ribonucleoprotein Prp4 | PRPF4 | * | 1.45 | x |
| **Q99878** | Histone H2A type 1-J | HIST1H2AJ | * | 0.73 | x |
| **P51809** | Vesicle-associated membrane protein 7 | VAMP7 | * | 1.88 | Membrane |
| **Q7L576** | Cytoplasmic FMR1-interacting protein 1 | CYFIP1 | * | – | Cytoplasm |

**Figure 3.**
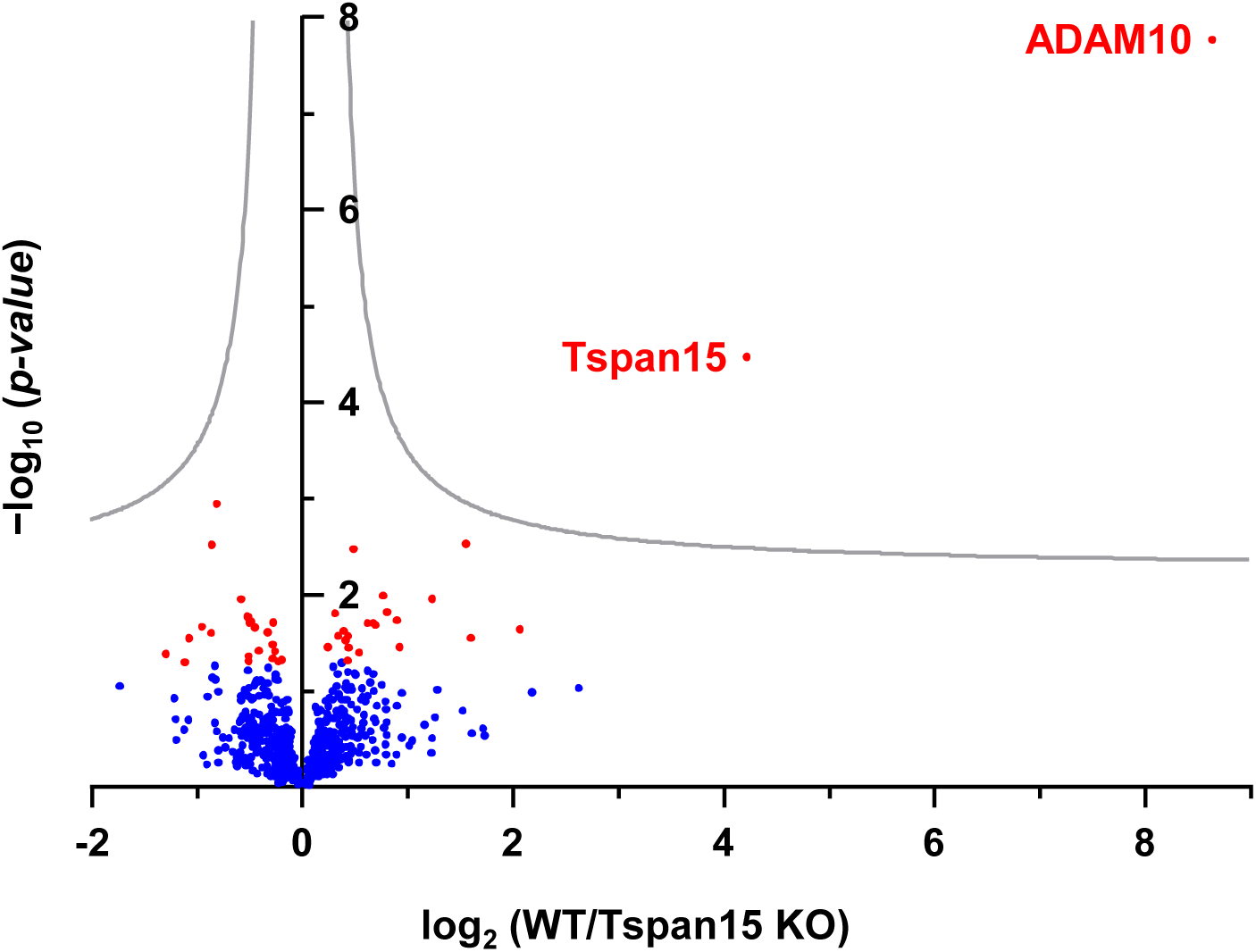
ADAM10 is the principal Tspan15-interacting protein in HEK-293T cells. Wildtype (WT) and Tspan15-knockout (KO) HEK-293T cells were lysed in 1% digitonin lysis buffer and immunoprecipitated with Tspan15 mAb 1C12 cross-linked to protein G sepharose beads. Proteins were identified by liquid chromatography coupled with tandem mass spectrometry (LC-MS/MS). Proteomic profiles of WT and Tspan15 KO HEK-293T immunoprecipitates are presented in a volcano plot to identify differentially expressed proteins. The minus log10 transformed *p*-value of each protein was plotted against the log2 transformed protein label free quantification ratio between the Tspan15 co-immunoprecipitation of WT samples and the control co-immunoprecipitation of Tspan15 KO samples. Proteins with significant fold change (*p*<0.05) are depicted in red; blue dots represent proteins with no significant changes in expression. A permutation-based false discovery rate estimation was applied and visualised as hyperbolic curves in grey.

### Tspan15 protein expression requires ADAM10

It is well established that TspanC8s promote the enzymatic maturation and trafficking of ADAM10 from the endoplasmic reticulum (ER) to the cell surface (Dornier *et al.* 2012; Haining *et al.* 2012; Prox *et al.* 2012). More recently, it was suggested that the reverse might be true, since ADAM10 knockdown reduced the trafficking of Tspan5 from the ER in HCT116 and U2OS cell lines (Saint-Pol *et al.* 2017). To investigate whether Tspan15 surface expression also requires ADAM10, flow cytometry was performed on CRISPR/Cas9 ADAM10-knockout cell lines. Tspan15 surface expression, as revealed by flow cytometry, was reduced by approximately 80% in the absence of ADAM10 in Jurkat, HEK-293T and A549 cells (Figure 4A). This observation held true in primary cells, since Tspan15 expression was almost undetectable following siRNA knockdown of ADAM10 in human umbilical vein endothelial cells (HUVECs) (Figure 4B). To determine whether the reduction in surface Tspan15 was consistent with a reduction in whole cell Tspan15, western blotting was performed on Tspan15 immunoprecipitates from Jurkat cell lysates. Similar to the flow cytometry data, Tspan15 protein expression was reduced by approximately 80% in the absence of ADAM10 (Figure 4C). Importantly, the loss of Tspan15 protein was not a consequence of reduced Tspan15 mRNA expression in the absence of ADAM10, as this was unaffected in ADAM10-knockout cell lines as measured by qRT-PCR (Figure 4D). These data demonstrate that ADAM10 and Tspan15 are each required for expression of the other, providing genetic evidence that the two proteins cooperate in a functional scissor complex.

**Figure 4.**
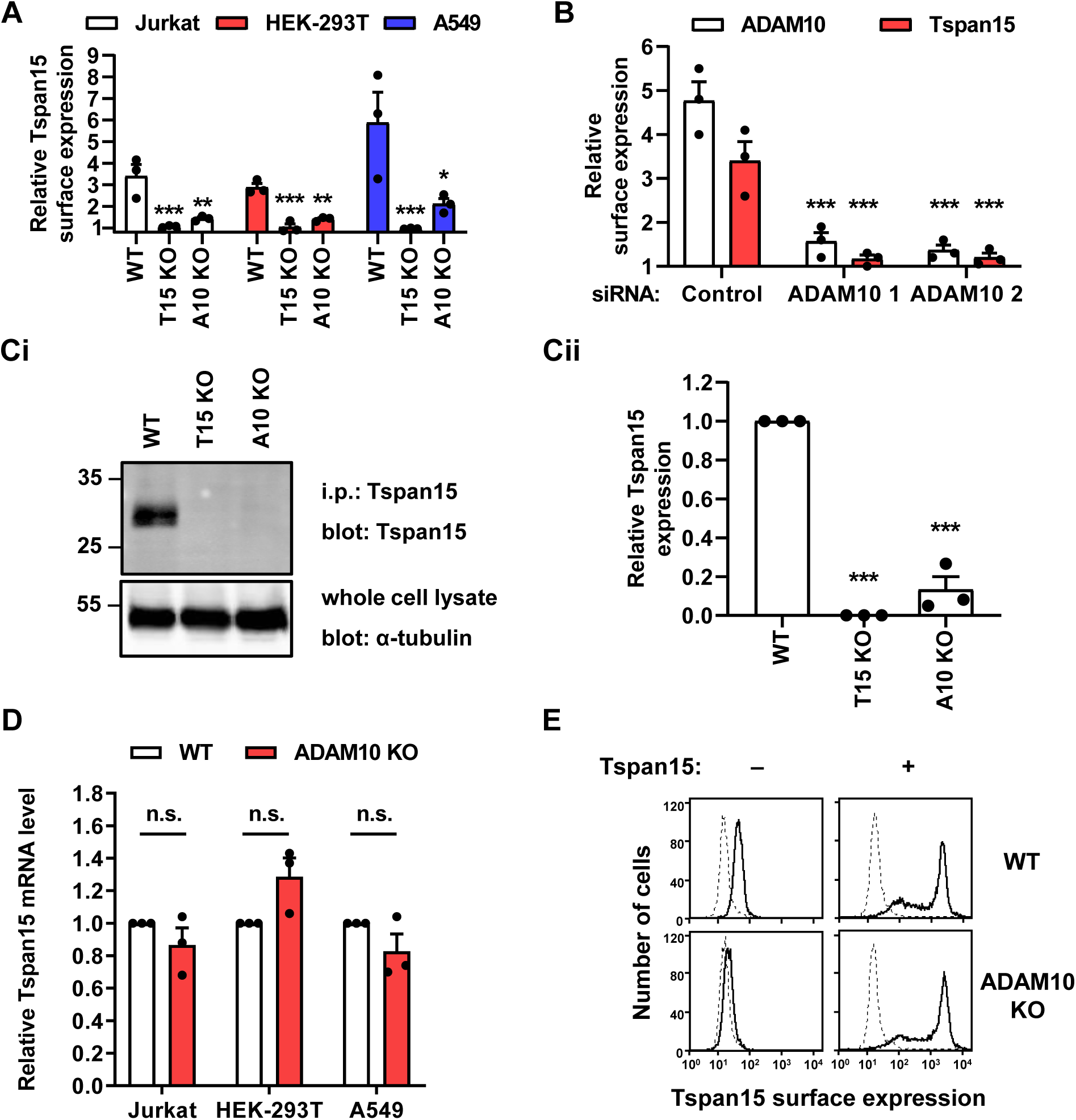
Tspan15 protein expression requires ADAM10. (A) Tspan15 surface expression in wildtype (WT), Tspan15-knockout (KO) and ADAM10 KO Jurkat, HEK-293T and A549 cell lines were analysed by flow cytometry with anti-Tspan15 mAb 1C12 or mouse IgG1 negative control antibody. Tspan15 surface expression is presented as the geometric mean fluorescence intensity of Tspan15 staining relative to the control staining. Error bars represent standard error of the mean from three independent experiments. Data were log-transformed and statistically analysed by a one-way ANOVA with Dunnett’s multiple comparisons test (\**p*<0.05, \*\**p*<0.01, \*\*\**p*<0.001 compared to WT). (B) HUVECs were transfected with two different ADAM10 siRNAs or a negative control siRNA and surface expression of ADAM10 and Tspan15 was measured by flow cytometry and analysed as described in panel A. (Ci) WT, Tspan15 KO and ADAM10 KO Jurkat cells were lysed in 1% Triton X-100 lysis buffer, immunoprecipitated with anti-Tspan15 mAb 5D4, and western blotted with the same antibody. Whole cell lysates were blotted with anti-α-tubulin mAb. (Cii) Tspan15 levels from panel Ci were quantitated and normalised to WT expression. Error bars represent standard errors of the mean from three independent experiments. Data were log-transformed and statistically analysed by a one-way ANOVA with a Tukey’s multiple comparisons test (\*\*\**p*<0.001 compared to WT). (D) Tspan15 mRNA level in WT and ADAM10 KO Jurkat, HEK-293T and A549 cells were assessed by qRT-PCR and presented relative to GAPDH housekeeping gene expression. Error bars represent standard errors of the mean from three independent experiments. Data were log-transformed and statistically analysed by a two-way ANOVA followed by Tukey’s multiple comparisons test (n.s., not significant). (E) WT and ADAM10 KO HEK-293T cells were transfected with empty vector control (−) or Tspan15 (+) expression constructs. Tspan15 surface expression was measured by flow cytometry as described in panel A. Histograms are representative of four independent experiments.

The requirement of ADAM10 for normal Tspan15 protein expression is inconsistent with the fact that human Tspan15 was successfully expressed in ADAM10-knockout MEFs to generate the immunogen for Tspan15 mAb generation (Figure 1A). To address whether over-expression of Tspan15 might overcome the requirement for ADAM10, HEK-293T cells were transiently transfected with Tspan15 and analysed by flow cytometry. In this over-expression scenario, Tspan15 expression was comparable in wild-type versus ADAM10-knockout cells (Figure 4E). Therefore, it appears that endogenous Tspan15 protein expression requires ADAM10, but over-expressed Tspan15 can bypass this requirement.

### The requirement of Tspan15 for ADAM10 surface expression is cell type dependent

To assess the extent to which Tspan15 is required for ADAM10 surface expression in the cells examined in Figure 4, flow cytometry for ADAM10 was performed. ADAM10 surface expression was reduced by approximately 60% and 70% in Tspan15-knockout HEK-293T and A549 cells, respectively, but was unaffected in Jurkat T cells (Figure 5A). In HUVECs, ADAM10 surface expression was reduced by approximately 40% following Tspan15 siRNA knockdown (Figure 5B). Therefore, the importance of Tspan15 for ADAM10 surface expression depends on the cell type.

**Figure 5.**
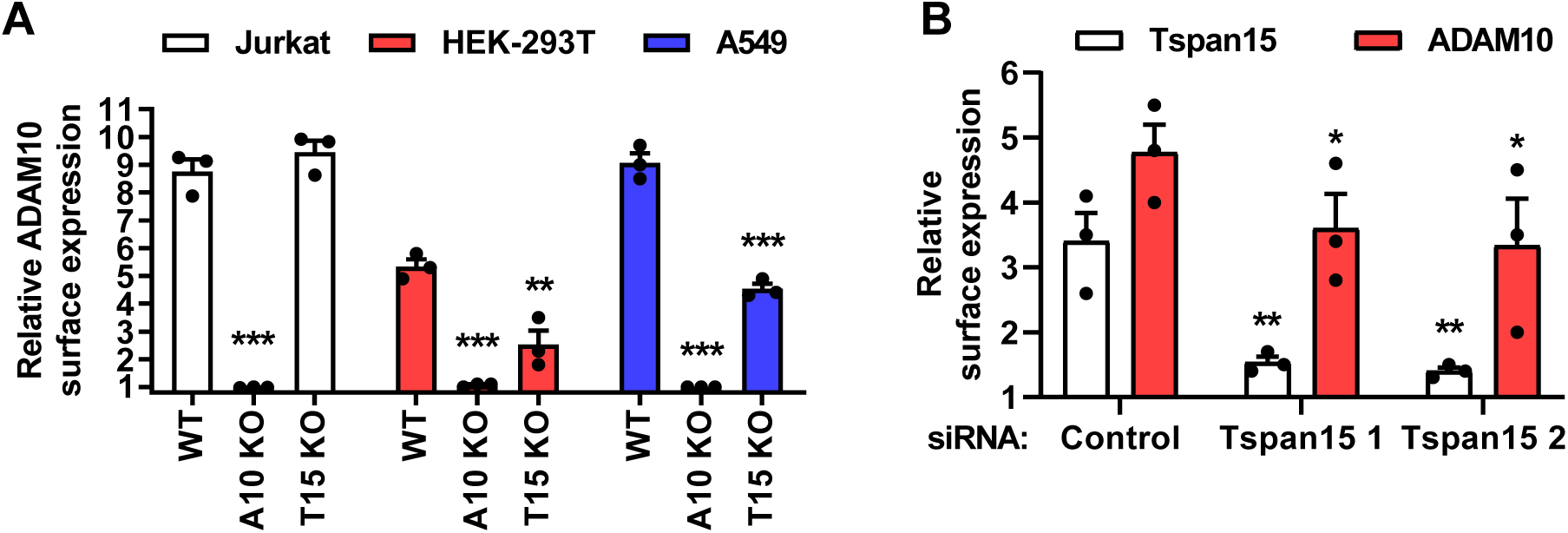
The requirement of Tspan15 for ADAM10 surface expression is cell type dependent. (A) ADAM10 surface expression in WT, ADAM10 KO and Tspan15 KO Jurkat, HEK-293T and A549 cells was measured by flow cytometry and quantitated as described in Figure 4A. (B) HUVECs were transfected with two different Tspan15 siRNAs or negative control siRNA and surface expression of ADAM10 was measured by flow cytometry and analysed as described in Figure 4A.

### ADAM10 and Tspan15 form dynamic bimolecular fluorescence complementation (BiFC) complexes

A single particle tracking study has previously reported an apparent diffusion coefficient for ADAM10 of 0.067 μm^2^/s in U2OS cells, which increased to 0.104 μm^2^/s following Tspan15 transfection (Jouannet *et al.* 2016). To more directly assess the lateral diffusion of ADAM10/Tspan15 complexes, and to image an ADAM10/Tspan15 dimer for the first time, bimolecular fluorescence complementation (BiFC) was employed. This technique allows formation of fluorescent protein dimers from two interacting proteins tagged with split fluorescent proteins, in this case Tspan15 fused to the N-terminal half of superfolder (sf)GFP and ADAM10 fused to the C-terminal half of sfGFP (Figure 6A). Formation of fluorescent BiFC ADAM10/Tspan15 dimers was confirmed by confocal microscopy of transfected HEK-293T cells, which showed a BiFC signal predominantly at the cell surface, similar to Tspan15 itself (Figure 6B). To examine the lateral diffusion of ADAM10/Tspan15 dimers on the apical membrane of transfected HEK-293T cells, fluorescence correlation spectroscopy (FCS) was used. The average number of fluorescent complexes/μm^2^ was 48 and the average diffusion speed was 0.19 μm^2^/s (Figure 6C-D). To then provide an indication of the oligomerisation state of the molecules, the average molecular brightness of ADAM10/Tspan15 dimers was determined using photon counting histogram (PCH) analysis. Of the FCS traces analysed, 58% fitted to a one-component PCH model (molecular brightness of 2.2 × 10^4^ cpm/s), whereas the remaining preferentially fitted to a two-component model that had dimmer (1.4×10^4^ cpm/s) and brighter (5.4×10^4^ cpm/s) subcomponents (Figure 6E). The molecular brightness of the dimmer traces was not significantly different from the one-component traces (Figure 6E). The presence of a brighter subcomponent suggests that some ADAM10/Tspan15 complexes were able to form larger clusters. Collectively, these data show that ADAM10 and Tspan15 can interact with each other to form dynamic BiFC dimers of varying stoichiometric ratio within nanoclusters.

**Figure 6.**
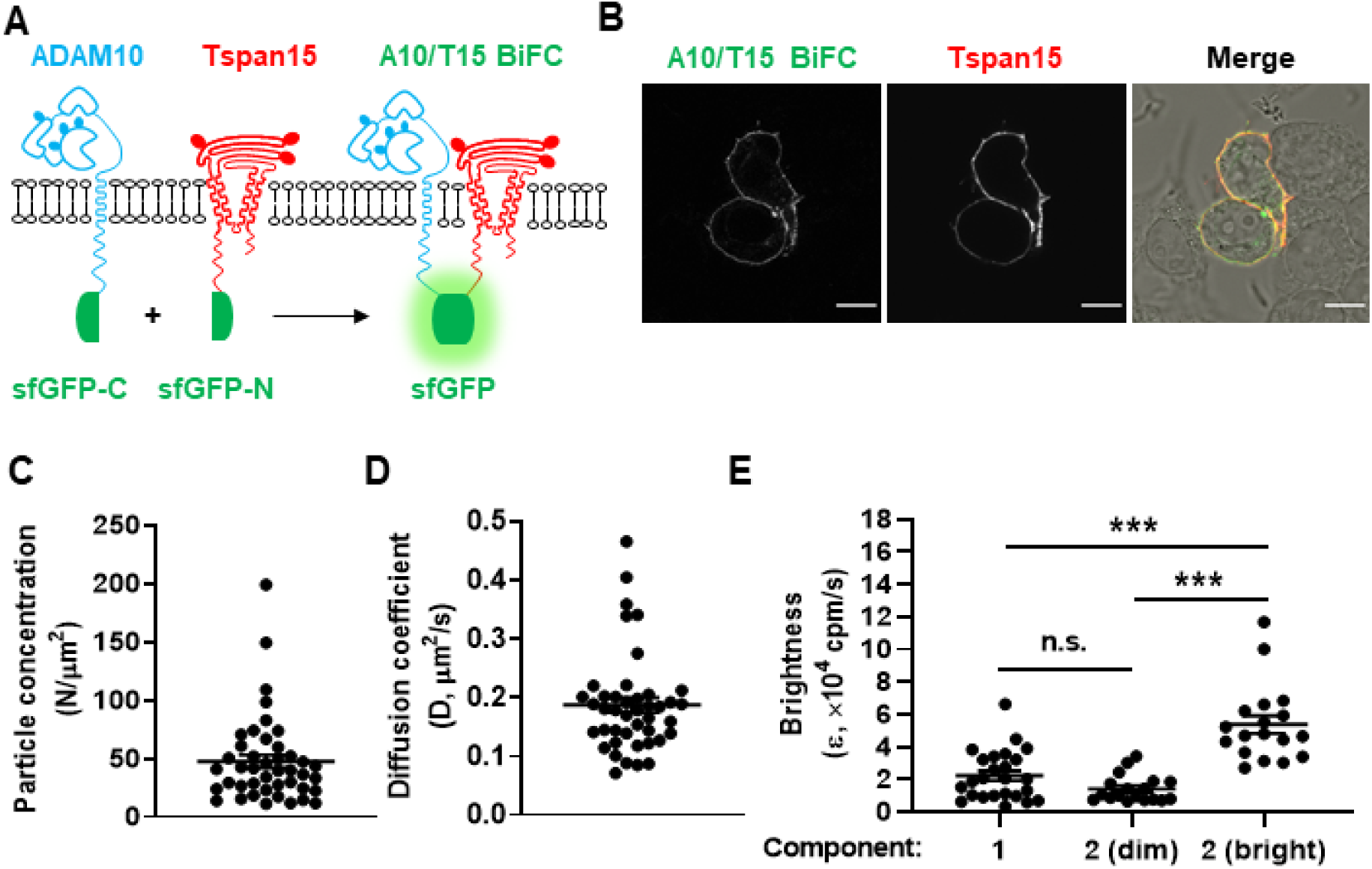
ADAM10 and Tspan15 form dynamic bimolecular fluorescence complementation (BiFC) complexes. (A) Schematic representation of ADAM10 tagged with the C-terminal half of superfolder GFP (sfGFP-C), Tspan15 tagged with the N-terminal half of superfolder GFP (sfGFP-N) and the predicted ADAM10/Tspan15 BiFC dimer. Solid ovals represent N-glycosylation. (B) HEK-293T cells were transfected with the ADAM10 and Tspan15 BiFC expression constructs, fixed and stained with Alexa Fluor® 647-conjugated Tspan15 mAb 5D4, and analysed by confocal microscopy. The image shown is representative of middle plane sections taken from two independent experiments (scale bar 10 µm). (C-D) Fluorescence correlation spectroscopy (FCS) measurements from the upper membrane of HEK-293T expressing the ADAM10/Tspan15 BiFC complexes were used to determine the average particle concentration (C) and diffusion co-efficient (D) of the complexes. (E) Fluorescence fluctuations from the FCS reads were also subjected to photon counting histogram (PCH) analysis to obtain the average molecular brightness (ε) of particles within the confocal volume. The FCS data were separated into groups that preferentially fit to a one-component or a two-component PCH model with dimmer and brighter subcomponents. Data were obtained from 43 individual measurements from three independent experiments. Error bars represent standard errors of the mean, N is the number of particles, and cpm is the counts per molecule. Data were log-transformed and statistically analysed by a one-way ANOVA followed by Tukey’s multiple comparisons test (\*\*\**p*<0.001).

### A synthetic ADAM10/Tspan15 fusion protein is a functional scissor

If ADAM10 and Tspan15 cooperate in a functional scissor complex, one prediction is that covalently linking the two proteins into one synthetic fusion protein (Figure 7A) would not disrupt subcellular localisation or scissor function. An ADAM10/Tspan15 fusion expression construct was generated and transfected into ADAM10/Tspan15 double knockout HEK-293T cells. Western blotting with ADAM10 and Tspan15 mAbs verified that the ADAM10/Tspan15 fusion protein had the predicted molecular weight of approximately 95 kD (Figure 7B), and flow cytometry (Figure 7C) and confocal microscopy (Figure 7D) showed that it was expressed at the cell surface. To test scissor function, ADAM10/Tspan15 double knockout HEK-293T cells were transfected with an expression construct for alkaline phosphatase-tagged betacellulin, a known ADAM10 substrate (Sahin *et al.* 2004), and shedding quantitated by measuring alkaline phosphatase activity released into the culture supernatant versus the non-shed activity remaining with the cells. Transfection of ADAM10/Tspan15 double knockout cells with the ADAM10/Tspan15 fusion protein construct restored betacellulin shedding to a level comparable to that of ADAM10 and Tspan15 transfected as individual constructs, in terms of both basal shedding and shedding induced by the ADAM10 activator N-ethylmaleimide (NEM) (Figure 7Ei). Betacellulin shedding was partially dependent on Tspan15, because ADAM10 transfection alone was not sufficient to fully restore shedding (Figure 7Ei). To confirm expression of the constructs used in these experiments, ADAM10 flow cytometry was used; the fusion protein was expressed at approximately two-fold greater levels than ADAM10 individually with Tspan15 (Figure 7Eii). These fusion protein data provide further evidence that ADAM10/Tspan15 exists as a functional scissor complex, because expression and function of the fusion protein are similar to when ADAM10 and Tspan15 are individually co-expressed.

**Figure 7.**
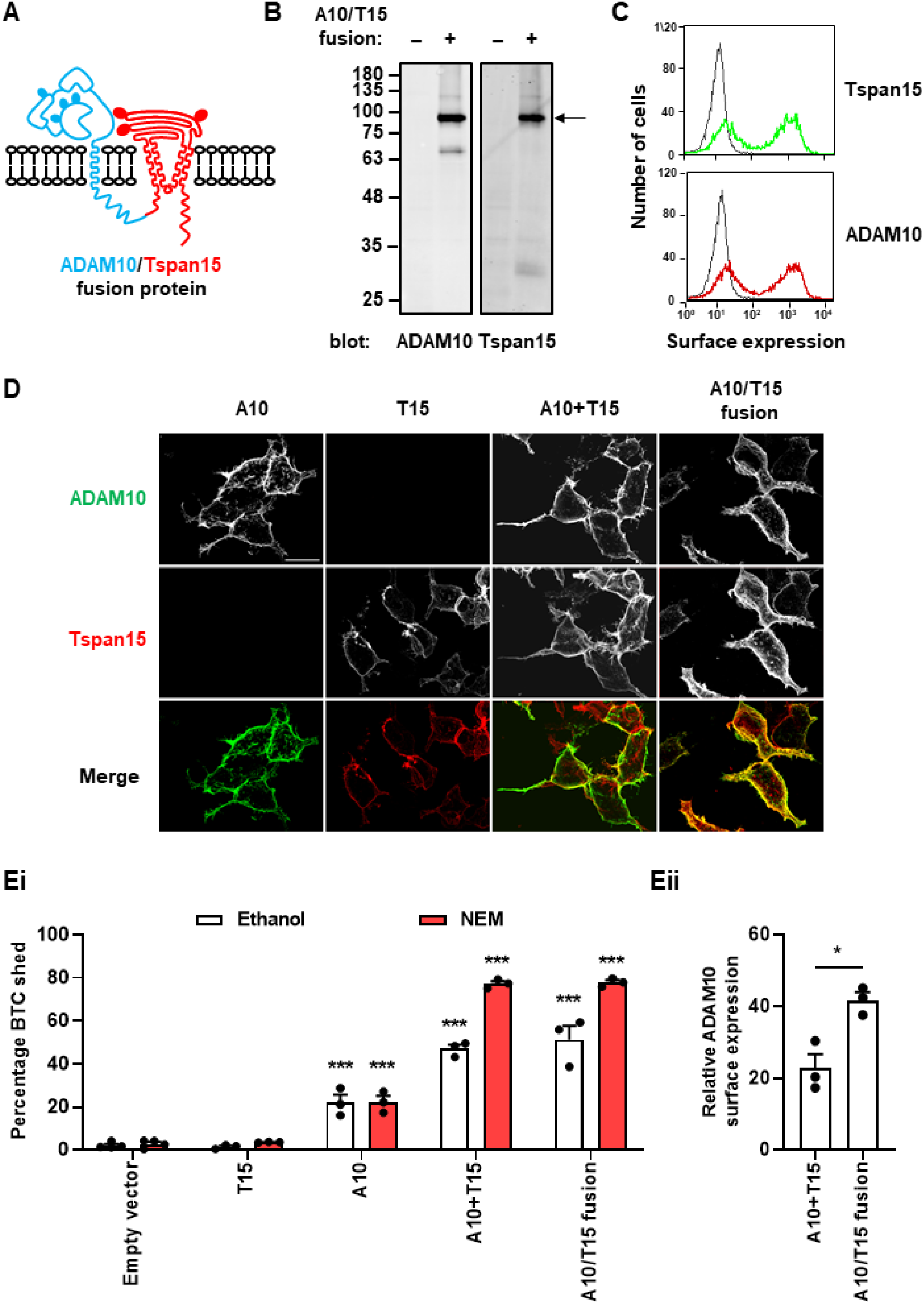
A synthetic ADAM10/Tspan15 fusion protein is a functional scissor. (A) Schematic representation of the synthetic ADAM10/Tspan15 fusion protein that has the C-terminus of ADAM10 physically linked to the N-terminus of Tspan15. Solid ovals represent N-glycosylation. (B) ADAM10/Tspan15 double KO HEK-293T cells were transfected with ADAM10/Tspan15 fusion construct, lysed in 1% digitonin lysis buffer in the presence of 10 µM ADAM10 inhibitor GI254023X, to prevent post-lysis auto-proteolysis, and western blotted for ADAM10 and Tspan15. (C) Cells described in panel B were assessed for surface expression of Tspan15 (green) and ADAM10 (red) by flow cytometry. Black traces represent isotype control staining. (D) ADAM10/Tspan15 double KO HEK-293T cells were transfected with the indicated expression constructs and analysed by confocal microscopy using anti-ADAM10 (green) and Tspan15 (red) mAbs. The cells were non-permeabilised and images are maximum intensity projections of confocal z-stacks. The scale bar on the upper left image is 30 µm. (Ei) ADAM10/Tspan15 double KO HEK-293T cells were co-transfected with alkaline phosphatase-tagged betacellulin (BTC) and Tspan15, ADAM10, ADAM10 and Tspan15, the ADAM10/Tspan15 fusion, or an empty vector control. Cells were stimulated with 2 mM NEM or vehicle control and alkaline phosphatase activity was measured in the supernatant and whole cell lysates to quantitate the percentage of BTC shed. Data were arcsine-transformed and statistically analysed with a two-way ANOVA followed by a Tukey’s multiple comparisons test (\*\*\**p*<0.001 compared to empty vector-transfected control). (Eii) Transfected cells were assessed for surface expression of ADAM10 by flow cytometry and the data quantitated and analysed using a paired t-test (\**p*<0.05).

## DISCUSSION

This study reports the generation of the first mAbs to tetraspanin Tspan15, which is one of six TspanC8 tetraspanins that regulate the molecular scissor ADAM10 (Matthews *et al.* 2017; Matthews *et al.* 2017; Saint-Pol *et al.* 2017) and was recently shown to promote cancer (Zhang *et al.* 2018). These mAbs enabled three major findings: (1) ADAM10 is the principal Tspan15-interacting protein; (2) Tspan15 expression requires ADAM10; and (3) a synthetic ADAM10/Tspan15 fusion protein is a functional scissor. These data strongly suggest that ADAM10 and Tspan15 exist as a heterodimeric scissor complex.

The discovery that Tspan15 expression requires ADAM10 provides genetic evidence that the principal role of Tspan15 is to regulate ADAM10. This finding is consistent with a recent study on Tspan5 and ADAM10 (Saint-Pol *et al.* 2017). In that study, an approximate 70% siRNA knockdown of ADAM10 in the HCT116 and U2OS cell lines reduced Tspan5 expression by 40%, and ADAM10 was shown to be important for trafficking of Tspan5 out of the ER (Saint-Pol *et al.* 2017). In the current study, we were able to investigate Tspan15 expression in the complete absence of ADAM10 by CRISPR/Cas9 knockout in three different cell lines, namely A549, HEK-293T and Jurkat. In each ADAM10-knockout cell line, Tspan15 expression was reduced by approximately 80%, as measured by flow cytometry and western blotting. Primary HUVECs yielded similar data, since 85-90% ADAM10 knockdown resulted in an equivalent reduction in Tspan15 protein expression. Importantly, Tspan15 mRNA expression levels were unaffected by ADAM10 knockout, indicating a specific effect on Tspan15 protein. Thus, Tspan15 and ADAM10 appear to be each required for the normal expression of the other. Indeed, it is well established that TspanC8s are important for ADAM10 maturation and trafficking to the cell surface (Dornier *et al.* 2012; Haining *et al.* 2012; Prox *et al.* 2012; Zhou *et al.* 2014; Jouannet *et al.* 2016; Noy *et al.* 2016; Virreira Winter *et al.* 2016; Reyat *et al.* 2017; Brummer *et al.* 2018; Seipold *et al.* 2018; Shah *et al.* 2018; Brummer *et al.* 2019), and we extended these observations by showing that ADAM10 surface expression was partially reduced in the absence of Tspan15 in A549, HEK-293T and HUVECs. Expression of other TspanC8s by these cells (Haining *et al.* 2012; Matthews *et al.* 2017) is likely to be the reason that the reduction is only partial. However, the effect of Tspan15 on ADAM10 expression is cell type dependent, because no ADAM10 reduction was observed in Tspan15-knockout Jurkat T cells. It remains unclear why this is the case, but it is possible that Tspan15 protein expression is low in these cells, relative to other tetraspanins.

The previously discussed finding that Tspan15 expression requires ADAM10 suggests that the two proteins might exist as a functional complex. A prediction of this idea is that physically linking the two proteins into a single fusion protein would yield a functional scissor protein. Indeed, we found that an ADAM10/Tspan15 fusion protein was expressed at the cell surface at the predicted molecular weight, and was functional in a betacellulin shedding assay in a manner that was comparable to when the two proteins were expressed individually. In this shedding assay in the HEK-293T cell line, betacellulin shedding was entirely dependent on ADAM10, as was previously reported in MEFs (Sahin *et al.* 2004). Our ADAM10/Tspan15 fusion protein is the first example of physically linking a tetraspanin to its partner protein, as a means of demonstrating that the two proteins can function as a complex. This approach may be useful for the investigation of other tetraspanins and their partners. Moreover, it may facilitate attempts to generate the first high-resolution structure of a tetraspanin/partner protein complex, which could provide fundamental new insights comparable to the recently-reported first crystal structure of a tetraspanin (Zimmerman *et al.* 2016). To date, the only structural study on a tetraspanin with its partner is the relatively low resolution (6 Å) cryo-electron microscopy study of tetraspanin uroplakins Ia and Ib with their uroplakin partners II and IIIa (Min *et al.* 2006).

Our study represents the first use of BiFC to enable the fluorescent imaging of a tetraspanin with its partner. BiFC uses two fusion proteins containing each half of a fluorescent protein, such as GFP, linked to the two proteins of interest, in this case ADAM10 and Tspan15. If the two proteins interact, the two halves of GFP will fold to form a fluorescent GFP molecule, and therefore any fluorescent signal can only be the result of a successful ADAM10/Tspan15 heterodimer. Consistent with the idea that ADAM10 and Tspan15 exist as a functional complex, ADAM10/Tspan15 BiFC dimers were formed and localised predominantly to the plasma membrane, similar to the localisation of Tspan15 itself. To investigate the dynamics of ADAM10/Tspan15 dimers for the first time in living cells, FCS was used to investigate its diffusion speed and clustering in the plasma membrane of transfected HEK-293T cells. An average diffusion co-efficient of 0.19 μm^2^/s was similar to that previously reported for ADAM10 (0.10 μm^2^/s) on Tspan15-transfected U2OS cells (Jouannet *et al.* 2016). However, that study measured ADAM10 rather than ADAM10/Tspan15 dimers (Jouannet *et al.* 2016), therefore a proportion of the diffusion speed will have been contributed by other ADAM10/TspanC8 complexes. We also used FCS to assess the molecular brightness of ADAM10/Tspan15 BiFC dimers, to estimate the degree of clustering. The existence of some traces containing a brighter Tspan15/ADAM10 component was indicative of larger cluster formation, as has been observed for TspanC8s and other tetraspanins by super-resolution microscopy (Zuidscherwoude *et al.* 2015; Marjon *et al.* 2016; Dahmane *et al.* 2019). We propose that FCS imaging of BiFC dimers will be useful for the study of other tetraspanin/partner protein complexes.

One prediction of the idea that ADAM10 and Tspan15 exist as a heterodimeric complex is that ADAM10 would be the major Tspan15-interacting partner. Using mass spectrometry to identify proteins in Tspan15 immunoprecipitates from HEK-293T cells lysed in the relatively stringent 1% digitonin, ADAM10 was strikingly the most abundant protein. Indeed, ADAM10 was almost 400 times more abundant in Tspan15 immunoprecipitates from wild-type cells versus control Tspan15-knockout cells, and the only other protein identified above the false discovery rate threshold was Tspan15 itself. These data are consistent with two previous studies in which GFP-tagged Tspan15 was expressed in U2OS or HepG2 cells, which were lysed in relatively non-stringent 1% Brij97 and Tspan15 immunoprecipitated via the GFP tag (Jouannet *et al.* 2016; Sidahmed-Adrar *et al.* 2019). For example, mass spectrometry identified ADAM10 as 15-fold more abundant than any other Tspan15-interacting proteins (Jouannet *et al.* 2016). However, interpretation was complicated in these two studies because of the use of less stringent detergent and the resulting large number of tetraspanin-associated proteins identified, the majority of which are likely to be indirectly associated with Tspan15. In our study, in addition to ADAM10, 26 proteins were identified as being significantly more detected in wild-type versus Tspan15-knockout samples, but the most differential of these was only four-fold higher in wild-type. Thus, it remains to be determined whether any of these are definitive ADAM10/Tspan15-interacting proteins that impact functionally on the complex. Of note, we did not identify beta-transducin repeat containing E3 ubiquitin protein ligase (BTRC) amongst the putative Tspan15-interacting proteins. BTRC was recently reported to co-immunoprecipitate with Tspan15 from lysates of oesophageal squamous cell carcinoma cell lines, and was implicated in promotion of metastasis by Tspan15 via the NF-κB pathway (Zhang *et al.* 2018). The Human Protein Atlas (www.proteinatlas.org) indicates that BTRC mRNA has a broad expression profile in cell lines, including HEK-293T (Thul *et al.* 2017), suggesting that we should have identified BTRC if it is a *bone fide* Tspan15-interacting protein. The findings of our study lead us to speculate that Tspan15 may be promoting cancer via ADAM10, most likely due to cleaving of specific substrates that enhance the cancer phenotype. Interestingly, gene expression data suggests that Tspan15 is strongly upregulated in cholangiocarcinoma and pancreatic adenocarcinoma (Tang *et al.* 2019), and is a marker of poor prognosis in pancreatic, renal and liver cancer (Uhlen *et al.* 2017). Moreover, Tspan15 has been recently implicated in oral squamous cell carcinoma (Hiroshima *et al.* 2019) and hepatocellular carcinoma (Sidahmed-Adrar *et al.* 2019).

The generation of mAbs to certain tetraspanins is a recognised difficulty in the tetraspanin field; there are no mAbs to many of the 33 human tetraspanins. To initiate this study, we generated the first Tspan15 mAbs using a novel strategy. Mice were immunised with ADAM10-knockout MEFs stably over-expressing human Tspan15, with the aim of focussing the mAb response on Tspan15 and preventing the larger ADAM10 from masking epitopes or preventing mAb binding by steric hindrance. Screening of hybridomas yielded four mouse anti-human Tspan15 mAbs, each recognising a similar epitope on the large extracellular loop of Tspan15. However, the importance of using ADAM10-knockout cells as the immunogen remains unclear. The immunisation and hybridoma generation was outsourced to Abpro, whose proprietary mouse may have contributed to the success of our approach. Furthermore, Tspan15 may be a relatively easy target, since eight of the 121 amino acids in the large extracellular loop are different between human and mouse, yielding 93% identity and the likelihood of distinct epitopes for mAb recognition. Most other TspanC8s are more conserved between human and mouse, with identities of 100, 77, 98, 96 and 97% for Tspan5, 10, 14, 17 and 33, respectively. A recent study overcame the problem of the identical human and mouse Tspan5 protein sequences by generating Tspan5 mAbs from Tspan5-knockout mice immunised with GFP-Tspan5-transfected cells and immunoprecipitates (Saint-Pol *et al.* 2017). Nevertheless, we hypothesise that our new method of using tetraspanin-transfected cells, with the tetraspanin partner(s) deleted, may be an effective strategy for future tetraspanin mAb generation.

To conclude, our findings strongly suggest that Tspan15 exists as an ADAM10/Tspan15 scissor complex, which may be a general theme for the other five TspanC8s. We therefore propose that ADAM10 be studied in the context of its associated TspanC8. This has implications for future work that should aim to determine how ADAM10/TspanC8 complexes are triggered to cleave specific substrates, and whether ADAM10/TspanC8 complexes can be therapeutically targeted to treat human disease.

## MATERIALS AND METHODS

### Antibodies

Primary antibodies were rabbit anti-FLAG (Sigma-Aldrich, Poole, UK), mouse or rabbit anti-α-tubulin (Cell Signaling Technology, London, UK), mouse anti-human ADAM10 11G2 (Arduise *et al.* 2008), mouse anti-Tspan5 TS52 (Saint-Pol *et al.* 2017) and negative control mouse IgG1 MOPC-21 (MP Biomedicals, Santa Ana, CA).

### Expression constructs

pLVX-EF1α-IRES-Puro FLAG-tagged human Tspan15 was generated by subcloning the FLAG-Tspan15 sequence into the pLVX-EF1α-IRES-Puro lentiviral plasmid (Clontech, Mountain View, CA). N-terminal FLAG-tagged human and mouse TspanC8s in the pEF6-*myc*-His plasmid (Invitrogen, Carlsbad, CA) were as described (Haining *et al.* 2012). A series of human/mouse Tspan15 chimeric constructs were generated using a two-step PCR method to incorporate mouse residues into the human Tspan15 large extracellular loop and *vice versa*. GFP-tagged human Tspan5/Tspan15 chimeras in the pEGFP-N1 plasmid were as described (Saint-Pol *et al.* 2017). Human Tspan15 tagged at the C-terminus with the N-terminal half of superfolder GFP was generated by subcloning human Tspan15 into a pcDNA3.1/zeo split superfolder GFP vector (Kilpatrick *et al.* 2012). Mouse ADAM10 tagged at the C-terminus with the C-terminal half of superfolder GFP was generated using a two-step PCR approach in which the GFP tag was subcloned into pcDNA3.1 mouse ADAM10 (Maretzky *et al.* 2005). pRK5M human ADAM10 was a gift from Rik Derynck (Addgene plasmid # 31717) (Liu *et al.* 2009). The pRK5M ADAM10-Tspan15 fusion construct was generated by subcloning human Tspan15 into pRK5M human ADAM10, at the C-terminus of ADAM10 to replace the myc tag. The alkaline phosphatase-tagged human betacellulin construct was kindly provided by Shigeki Higashiyama and Carl Blobel (Sahin *et al.* 2004).

### Cell culture and transfections

ADAM10-knockout mouse embryonic fibroblasts (MEFs) (Reiss *et al.* 2005), human embryonic kidney (HEK)-293T (HEK-293 cells expressing the large T-antigen of simian virus 40) and the human lung epithelial cell line A549 were cultured in DMEM (Sigma-Aldrich), and the human T cell line Jurkat was cultured in RPMI 1640 (Sigma-Aldrich), each supplemented with 10% fetal bovine serum (Thermo Fisher Scientific, Loughborough, UK), 4 mM L-glutamine,100 units/ml penicillin and 100 μg/ml streptomycin (Sigma-Aldrich). Human umbilical vein endothelial cells (HUVECs) were isolated from umbilical cords obtained with consent from the Human Biomaterials Resource Centre at the University of Birmingham, UK, and cultured in M199 supplemented with 10% fetal bovine serum, 4 mM glutamine, 90 μg/ml heparin (Sigma-Aldrich) and bovine brain extract (Wilson *et al.* 2014), and used between passages 3 and 6. HEK-293T cells were transfected using polyethyleneimine as described (Ehrhardt *et al.* 2006). A549 cells were transfected with Lipofectamine 2000 (Thermo Fisher Scientific). Jurkat cells were transfected by electroporation as previously described (Tomlinson *et al.* 2007). For siRNA knockdowns, HUVECs were transfected with 10 nM Silencer Select siRNA duplexes (Thermo Fisher Scientific) using Lipofectamine RNAiMAX (Thermo Fisher Scientific).

### Generation of CRISPR/Cas9-knockout cell lines

Two guide RNA sequences were selected for each of human ADAM10 and Tspan15 using the Wellcome Trust Sanger Institute’s CRISPR Finder tool (Hodgkins *et al.* 2015). The following primer pairs were used to encode these sequences: ADAM10 guide 1 (5’-CACCGCGTCTAGATTTCCATGCCCA-3’ and 5’-AAACTGGGCATGGAAATCTAGACGC-3’); ADAM10 guide 2 (5’-CACCGATACCTCTCATATTTACAC-3’ and 5’-

AAACGTGTAAATATGAGAGGTATC-3’); Tspan15 guide 1 (5’-CACCGGCGCGCGCTTCTCCTACCTC-3’ and 5’-AAACGAGGTAGGAGAAGCGCGCGCC-3’); Tspan15 guide 2 (5’-CACCGAGCGCCCAGGATGCCGCGCG-3’ and 5’-AAACCGCGCGGCATCCTGGGCGCTC-3’). Each primer pair was annealed and cloned into the pSpCas9 (BB)-2A-Puro (PX459) plasmid (a gift from Feng Zhang, Addgene plasmid #62988) (Ran *et al.* 2013). Cells were transfected with either of the guide constructs, clonal transfectants selected using 1, 2.5 and 0.5 μg/ml puromycin (Thermo Fisher Scientific) for A549, HEK-293T and Jurkat cells, respectively, and knockouts confirmed by flow cytometry.

### Human platelet preparation

Washed human platelets were isolated from whole blood by centrifugation as described previously (McCarty *et al.* 2006). Blood samples were taken with consent from healthy donors.

### Generation of new Tspan15 mAbs

To create the immunogen for mouse anti-human Tspan15 mAb generation, ADAM10-knockout MEFs were transduced with lentivirus packaged with FLAG-tagged human Tspan15, produced in HEK-293T cells using a previously described protocol (Reyat *et al.* 2017). Clonal cells stably overexpressing Tspan15 were selected with 1 μg/ml puromycin and validated by anti-FLAG western blotting. Immunisation of mice and generation of hybridomas were outsourced to Abpro Therapeutics (Boston, MA).

### Antibody conjugation

Purified Tspan15 mAbs were concentrated to 1 mg/ml using Amicon Ultra Centrifugal Filters (Merck, Watford, UK) and conjugated to Alexa Fluor® 647 fluorophore with a succinimidyl ester reactive dye (Thermo Fisher Scientific) at an antibody-to-dye mass ratio of 10:1 for one hour at room temperature. Excess unbound dye was removed using Zeba Spin Desalting Columns (Thermo Fisher Scientific). The concentration and degree of labelling of the conjugated mAbs were measured spectrophotometrically.

### Flow cytometry

2.5–5 × 10^5^ cells were stained with primary antibodies at 10 μg/ml, followed by fluorescein isothiocyanate (FITC)-conjugated sheep anti-mouse (Sigma-Aldrich) or allophycocyanin (APC)-conjugated goat anti-mouse (Thermo Fisher Scientific) secondary antibodies. Samples were analysed on a FACSCalibur flow cytometer (BD Biosciences, Oxford, UK). Surface staining was quantitated by its geometric mean fluorescence intensity and presented as a ratio to the isotype control.

### Antibody binding competition assay

A549 cells were incubated with 10 µg/ml MOPC-21 or Tspan15 mAbs 1C12, 4A4, 5D4 or 5F4 for 30 minutes on ice. The cells were then stained with 10 µg/ml of each of the Alexa Fluor® 647-conjugated Tspan15 mAbs for 30 minutes on ice. Relative antibody binding was analysed by flow cytometry.

### Immunoprecipitation and western blotting

Immunoprecipitation and western blotting were performed as previously described (Noy *et al.* 2016). Western blotting used IRDye 680RD or 800CW-conjugated secondary antibodies (LI-COR Biosciences, Cambridge, UK) and an Odyssey Quantitative Infrared Imaging System (LI-COR Biosciences).

### Sample preparation for mass spectrometry

2 × 10^8^ wild-type or Tspan15-knockout HEK-293T cells were lysed in 1% digitonin lysis buffer containing protease inhibitor cocktail (Sigma-Aldrich). Lysates were pre-cleared with protein G sepharose prior to immunoprecipitation with Tspan15 mAb 1C12 chemically cross-linked to protein G sepharose with dimethyl pimelimidate (Thermo Fisher Scientific). Five independent immunoprecipitations were carried out for each cell type. Immunoprecipitation samples in non-reducing Laemmli buffer were subjected to a modified single-pot solid-phase-enhanced sample preparation (SP3) protocol (Hughes *et al.* 2019). Briefly, 10 µl of a 4 µg/µl bead slurry of Sera-Mag SpeedBeads A and B (GE Healthcare, Chicago, IL) were added to the samples. Protein binding to the magnetic beads was achieved by adding acetonitrile to a final volume of 70% (v/v) and mixing at 1200 rpm at 24 °C for 30 minutes in a Thermomixer (Eppendorf, Hamburg, Germany). Magnetic beads were retained in a DynaMag-2 magnetic rack (Thermo Fisher Scientific) and the supernatant was discarded. Disulfide bridges were reduced by adding 20 µl of 30 mM dithiothreitol (Biozol, Eching, Germany) and incubating at 1200 rpm at 37 °C for 30 minutes. Cysteines were alkylated by adding 25 µl of 80 mM iodoactemamide (Sigma-Aldrich) and incubating at 1200 rpm at 24 °C for 30 minutes in the dark in a Thermomixer. The reaction was quenched by adding 3 µl of 200 mM dithiothreitol. Protein binding to the beads was repeated in 70% (v/v) acetonitrile for 30 minutes. After removing the solvent, beads were washed twice in 200 µl 70% (v/v) ethanol and twice in 180 µl of 100% (v/v) acetonitrile. Next, 250 ng of LysC and 250 ng of trypsin (Promega, Mannheim, Germany) were added in 20 µl of 50 mM ammonium bicarbonate (Sigma Aldrich). The protein digestion was performed for 16 hours at room temperature. Samples were acidified with formic acid to a final concentration of 1% (v/v) and placed in a magnetic rack. The supernatants were transferred into fresh 0.5 ml protein LoBind tubes (Eppendorf). A volume of 20 µl of 2% (v/v) dimethyl sulfoxide was added to the beads and subjected to sonication for 30 seconds in a water bath. Tubes were placed in the magnetic rack and the supernatants were transferred. The samples were dried in a vacuum centrifuge and dissolved in 20 µl 0.1% formic acid.

### Liquid chromatography coupled with tandem mass spectrometry (LC-MS/MS) analysis

Samples were analyzed by LC-MS/MS for relative label free protein quantification. A volume of 10 µl per sample was separated on a nanoLC system (EASY-nLC 1200, Thermo Fisher Scientific) using an in-house packed C18 column (30 cm × 75 µm ID, ReproSil-Pur 120 C18-AQ, 1.9 µm; Dr. Maisch GmbH, Ammerbuch, Germany) with a binary gradient of water (A) and acetonitrile (B) containing 0.1% formic acid at 50 °C column temperature and a flow rate of 250 nl/minute (gradient: 0 minutes 2.4% B; 2 minutes, 4.8% B; 92 minutes, 24% B; 112 minutes, 35.2% B; 121 minutes, 60% B).

The nanoLC was coupled online via a nanospray flex ion source (Proxeon, Thermo Fisher Scientific) equipped with a PRSO-V2 column oven (Sonation, Biberach an der Riss, Germany) to a Q-Exactive HF mass spectrometer (Thermo Fisher Scientific). Full MS spectra were acquired at a resolution of 120,000. The top 15 peptide ions were chosen for Higher-energy C-trap Dissociation (HCD) (normalized collision energy of 26%, automatic gain control 1E+5 ions, intensity threshold 5E+3 ions, maximum ion trapping time 100 ms). Fragment ion spectra were acquired at a resolution of 15,000. A dynamic exclusion of 120 s was used for peptide fragmentation.

### Data analysis and label free quantification

The raw data was analyzed by Maxquant software (maxquant.org, Max-Planck Institute Munich) version 1.6.6.0 (Cox *et al.* 2014). The MS data was searched against a fasta database of *Homo sapiens* from UniProt (download: June 12^th^ 2019, 20962 entries). Trypsin was defined as the protease. Two missed cleavages were allowed for the database search. The option first search was used to recalibrate the peptide masses within a window of 20 ppm. For the main search, peptide and peptide fragment mass tolerances were set to 4.5 and 20 ppm, respectively. Carbamidomethylation of cysteine was defined as static modification. Acetylation of the protein N-terminus and oxidation of methionine were set as variable modifications. The false discovery rate for both peptides and proteins was adjusted to less than 1%. Label free quantification of proteins required at least two ratio counts of unique peptides. Only unique peptides were used for quantification. The option “match between runs” was enabled with a matching time of 1 minute.

The protein label free quantification intensities were log2 transformed and a two-sided Student’s *t*-test was applied to evaluate the significance of proteins with changed abundance between the TSPAN15 immunoprecipitation from wild-type samples and the control immunoprecipitation from Tspan15-knockout samples. Additionally, a permutation-based false discovery rate estimation was used (Tusher *et al.* 2001).

### Quantitative reverse-transcription PCR (qRT-PCR)

RNA was extracted using the RNeasy Mini kit (Qiagen, Manchester, UK) from cells homogenised with QIAshredder columns (Qiagen). Complementary DNA was generated using the High Capacity cDNA Reverse Transcription kit (Thermo Fisher Scientific) and subjected to TaqMan qPCR assays (Thermo Fisher Scientific) for Tspan15 (Hs00202548_m1) and GAPDH (Hs02758991_g1) and analysed as described (Reyat *et al.* 2017). All qPCR data were normalised to GAPDH as the internal loading control.

### Confocal microscopy

All reagents are from Sigma-Aldrich unless otherwise stated. Cells were fixed with 10% formalin for 15 minutes and washed three times with PBS (0.01 M phosphate, 0.0027 M KCl, 0.137 M NaCl, pH 7.4) before blocking in PBS containing 1% BSA, 2% goat serum (or mouse serum for direct staining with conjugated mouse primary antibodies) and 0.1% saponin for 20 minutes to permeabilise the cells. The same buffer was used for antibody dilutions. Cells were incubated with antibodies for one hour and washed three to four times with PBS after incubations. For direct labelling of Tspan15, cells were stained with 5 µg/ml Alexa Fluor® 647-conjugated Tspan15 mAb (5D4). For simultaneous staining of ADAM10 and Tspan15, cells were first stained with 0.5 µg/ml ADAM10 mAb (11G2) and then secondarily labelled with Alexa Fluor®-conjugated goat anti-mouse antibody, followed by direct staining of Tspan15 as described above. Confocal Z-stacks of stained cells were taken at 1 µm intervals on a Leica TCS SP2 confocal microscope (Leica Biosystems, Wetzlar, Germany) with 488 nm Argon and 633 nm Helium/Neon laser lines with a 63x 1.4 NA oil objective. For imaging of non-permeabilised cells, the Nikon A1R confocal system (Nikon, Tokyo, Japan), equipped with a 100x 1.4 NA oil objective and similar lasers to those above, was used for acquisition of z-stacks.

### Fluorescence correlation spectroscopy (FCS)

1.5 × 10^4^ HEK-293T cells were seeded onto Nunc Lab-Tek 8-well glass-bottomed chamber slides (0.13-0.17 mm thick, #1.0 borosilicate glass) (Thermo Fisher Scientific) pre-coated with 10 µg/ml poly-D-lysine (Sigma-Aldrich). 24 hours later, cells were transfected with ADAM10 and Tspan15 split superfolder GFP (sfGFP) expression constructs. After a further 24 hours, media was replaced with 200 μl/well HEPES-buffered saline solution (2 mM sodium pyruvate, 145 mM NaCl, 10 mM D-glucose, 5 mM KCl, 1 mM MgSO_4_.7H_2_O, 10 mM HEPES, 1.3 mM CaCl_2_, 1.5 mM NaHCO_3_, pH 7.45) and equilibrated for 10 minutes at 22 °C prior to FCS recording on a Zeiss LSM 510NLO Confocor 3 microscope (Carl Zeiss, Jena, Germany). Fluorescence fluctuations were collected using a c-Apochromat 40x 1.2 NA water immersion objective using 488 nm excitation with emission collected through a 505-600 long pass filter. FCS acquisition, autocorrelation and photon counting histogram (PCH) analyses were performed as described previously (Ayling *et al.* 2012). In brief, the confocal volume was calibrated with 20 nM rhodamine 6G dye on each experimental day. For cell measurements, the detection volume was positioned in x-y over the cell of interest using a live confocal image, and then on the apical cell membrane following a z-intensity scan using ~0.04 kW/cm^2^ laser power. Fluctuation traces were recorded at 22 °C on the apical membrane for each cell for 30 seconds using ~0.17 kW/cm^2^ laser power. Autocorrelation and PCH analyses were performed in Zen 2012 (Carl Zeiss), with the first 5 seconds of fluctuations routinely removed to adjust for photobleaching. Autocorrelation curves were fitted to a two-component, 2D diffusion model with a pre-exponential term to account for sfGFP blinking, with an offset added where necessary to obtain average dwell times and particle numbers. As previously demonstrated, component 1 (300-600 μs) represents a dark state of the sfGFP fluorophore, whilst component 2 represents the dwell time of the sfGFP-labelled complex (Ayling *et al.* 2012; Kilpatrick *et al.* 2012). Diffusion coefficients of the sfGFP-labelled complex were calculated for each trace using the equation D = ω_0_^2^/4.τ_D_, where ω_0_ is the radius of the detection volume (obtained from calibration) and τ_D_ the average dwell time of component 2. Particle numbers of component 2 were expressed as particles per μm^2^, calculated from N = N(τ_D2_)/ π.ω_0_^2^, where N(τ_D2_) is the fractional contribution of component 2 to the total particle number determined from the autocorrelation curve. Molecular brightness of complexes was determined using PCH analyses carried out on the same fluctuation traces as used for autocorrelation analyses using Zen 2012. Traces were binned at 100 μs and fitted to either a 1- or 2-component PCH model with the first order correction fixed at 0.3, as determined from the calibration read. Data are shown as values obtained from individual cell membranes, obtained over ‘n’ independent transfections.

### Betacellulin shedding assay

ADAM10/Tspan15 double knockout HEK-293T cells were transfected in 24-well plates with 200 ng of alkaline phosphatase-conjugated betacellulin expression construct and 50 ng total of ADAM10, Tspan15, ADAM10 plus Tspan15, or the ADAM10/Tspan15 fusion construct. 24 hours post transfection, cells were washed and stimulated with 2 mM N-ethylmaleimide (NEM) (Sigma-Aldrich), or ethanol vehicle control, for 2.5 hours in Opti-MEM reduced serum media (Thermo Fisher Scientific). Alkaline phosphatase activity in supernatant and cell lysate samples were measured using p-nitrophenyl phosphate (pNPP) substrate (Sigma-Aldrich) and a VICTOR X3 Multilabel Plate Reader (Perkin Elmer, Seer Green, UK). The supernatant activity as a percentage of the total was calculated to determine the percentage shedding.

### Statistical analyses

Relative data were normalised by either log- or arscine-transformation before being statistical analysed using ANOVA with post-hoc multiple comparison tests, as indicated in the figure legends.

## ACKNOWLEDGMENTS

We are grateful for members of the *Cells & Molecules* research theme in the School of Biosciences for their helpful comments on this project. We also thank the Birmingham Advanced Light Microscope Facility and COMPARE for advice on fluorescence microscopy. This work was funded by a British Heart Foundation PhD Studentship and COMPARE grant which supported C.Z.K. (FS/18/9/33388), a Biotechnology and Biological Sciences Research Council Project Grant which supported N.H. (BB/P00783X/1), a British Heart Foundation Project Grant (PG/13/92/30587) which supported P.J.N., and Biotechnology and Biological Sciences Research Council PhD Studentships which supported J.S. and A.L.M. This work was also supported by the Deutsche Forschungsgemeinschaft (German Research Foundation) within the framework of the Munich Cluster for Systems Neurology (EXC 2145 SyNergy) and by the BMBF through CLINSPECT-M to S.F.L. Further support was provided by a Deutsche Forschungsgemeninschaft grant (DFG-SFB877-A3) to P.S.

## COMPETING INTERESTS

The authors declare no competing interests.

## REFERENCES

Arduise, C., T. Abache, L. Li, M. Billard, A. Chabanon, A. Ludwig, P. Mauduit, C. Boucheix, E. Rubinstein and F. Le Naour (2008). Tetraspanins regulate ADAM10-mediated cleavage of TNF-alpha and epidermal growth factor. J Immunol 181: 7002–7013.

Ayling, L. J., S. J. Briddon, M. L. Halls, G. R. Hammond, L. Vaca, J. Pacheco, S. J. Hill and D.n M. Cooper (2012). Adenylyl cyclase AC8 directly controls its micro-environment by recruiting the actin cytoskeleton in a cholesterol-rich milieu. J Cell Sci 125: 869–886. 10.1242/jcs.091090.

Brummer, T., S. A. Muller, F. Pan-Montojo, F. Yoshida, A. Fellgiebel, T. Tomita, K. Endres and S. F. Lichtenthaler (2019). NrCAM is a marker for substrate-selective activation of ADAM10 in Alzheimer’s disease. EMBO Mol Med 11: e9695. 10.15252/emmm.201809695.

Brummer, T., M. Pigoni, A. Rossello, H. Wang, P. J. Noy, M. G. Tomlinson, C. P. Blobel and S. F. Lichtenthaler (2018). The metalloprotease ADAM10 (a disintegrin and metalloprotease 10) undergoes rapid, postlysis autocatalytic degradation. FASEB J 32: 3560–3573. 10.1096/fj.201700823RR.

Cox, J., M. Y. Hein, C. A. Luber, I. Paron, N. Nagaraj and M. Mann (2014). Accurate proteome-wide label-free quantification by delayed normalization and maximal peptide ratio extraction, termed MaxLFQ. Mol Cell Proteomics 13: 2513–2526. 10.1074/mcp.M113.031591.

Dahmane, S., C. Doucet, A. Le Gall, C. Chamontin, P. Dosset, F. Murcy, L. Fernandez, D. Salas, E. Rubinstein, M. Mougel, M. Nollmann and P. E. Milhiet (2019). Nanoscale organization of tetraspanins during HIV-1 budding by correlative dSTORM/AFM. Nanoscale 11: 6036–6044. 10.1039/c8nr07269h.

Dornier, E., F. Coumailleau, J. F. Ottavi, J. Moretti, C. Boucheix, P. Mauduit, F. Schweisguth and E. Rubinstein (2012). TspanC8 tetraspanins regulate ADAM10/Kuzbanian trafficking and promote Notch activation in flies and mammals. J Cell Biol 199: 481–496. 10.1083/jcb.201201133.

Ehrhardt, C., M. Schmolke, A. Matzke, A. Knoblauch, C. Will, V. Wixler and S. Ludwig (2006). Polyethylenimine, a cost-effective transfection reagent. Signal Transduction 6: 179–184.

Haining, E. J., J. Yang, R. L. Bailey, K. Khan, R. Collier, S. Tsai, S. P. Watson, J. Frampton, P. Garcia and M. G. Tomlinson (2012). The TspanC8 subgroup of tetraspanins interacts with A disintegrin and metalloprotease 10 (ADAM10) and regulates its maturation and cell surface expression. J Biol Chem 287: 39753–39765. 10.1074/jbc.M112.416503.

Hiroshima, K., M. Shiiba, N. Oka, F. Hayashi, S. Ishida, R. Fukushima, K. Koike, M. Iyoda, D. Nakashima, H. Tanzawa and K. Uzawa (2019). Tspan15 plays a crucial role in metastasis in oral squamous cell carcinoma. Exp Cell Res: 111622. 10.1016/j.yexcr.2019.111622.

Hodgkins, A., A. Farne, S. Perera, T. Grego, D. J. Parry-Smith, W. C. Skarnes and V. Iyer (2015). WGE: a CRISPR database for genome engineering. Bioinformatics 31: 3078–3080. 10.1093/bioinformatics/btv308.

Hughes, C. S., S. Moggridge, T. Muller, P. H. Sorensen, G. B. Morin and J. Krijgsveld (2019). Single-pot, solid-phase-enhanced sample preparation for proteomics experiments. Nat Protoc 14: 68–85. 10.1038/s41596-018-0082-x.

Jouannet, S., J. Saint-Pol, L. Fernandez, V. Nguyen, S. Charrin, C. Boucheix, C. Brou, P. E. Milhiet and E. Rubinstein (2016). TspanC8 tetraspanins differentially regulate the cleavage of ADAM10 substrates, Notch activation and ADAM10 membrane compartmentalization. Cell Mol Life Sci 73: 1895–1915. 10.1007/s00018-015-2111-z.

Kilpatrick, L. E., S. J. Briddon and N. D. Holliday (2012). Fluorescence correlation spectroscopy, combined with bimolecular fluorescence complementation, reveals the effects of beta-arrestin complexes and endocytic targeting on the membrane mobility of neuropeptide Y receptors. Biochim Biophys Acta 1823: 1068–1081. 10.1016/j.bbamcr.2012.03.002.

Lichtenthaler, S. F., M. K. Lemberg and R. Fluhrer (2018). Proteolytic ectodomain shedding of membrane proteins in mammals-hardware, concepts, and recent developments. EMBO J 37. 10.15252/embj.201899456.

Liu, C., P. Xu, S. Lamouille, J. Xu and R. Derynck (2009). TACE-mediated ectodomain shedding of the type I TGF-beta receptor downregulates TGF-beta signaling. Mol Cell 35: 26–36. 10.1016/j.molcel.2009.06.018.

Maretzky, T., M. Schulte, A. Ludwig, S. Rose-John, C. Blobel, D. Hartmann, P. Altevogt, P. Saftig and K. Reiss (2005). L1 is sequentially processed by two differently activated metalloproteases and presenilin/gamma-secretase and regulates neural cell adhesion, cell migration, and neurite outgrowth. Mol Cell Biol 25: 9040–9053. 10.1128/MCB.25.20.9040-9053.2005.

Marjon, K. D., C. M. Termini, K. L. Karlen, C. Saito-Reis, C. E. Soria, K. A. Lidke and J. M. Gillette (2016). Tetraspanin CD82 regulates bone marrow homing of acute myeloid leukemia by modulating the molecular organization of N-cadherin. Oncogene 35: 4132–4140. 10.1038/onc.2015.449.

Matthews, A. L., C. Z. Koo, J. Szyroka, N. Harrison, A. Kanhere and M. G. Tomlinson (2018). Regulation of Leukocytes by TspanC8 Tetraspanins and the “Molecular Scissor” ADAM10. Front Immunol 9: 1451. 10.3389/fimmu.2018.01451.

Matthews, A. L., P. J. Noy, J. S. Reyat and M. G. Tomlinson (2017). Regulation of A disintegrin and metalloproteinase (ADAM) family sheddases ADAM10 and ADAM17: The emerging role of tetraspanins and rhomboids. Platelets 28: 333–341. 10.1080/09537104.2016.1184751.

Matthews, A. L., J. Szyroka, R. Collier, P. J. Noy and M. G. Tomlinson (2017). Scissor sisters: regulation of ADAM10 by the TspanC8 tetraspanins. Biochem Soc Trans 45: 719–730. 10.1042/BST20160290.

McCarty, O. J., S. D. Calaminus, M. C. Berndt, L. M. Machesky and S. P. Watson (2006). von Willebrand factor mediates platelet spreading through glycoprotein Ib and alpha(IIb)beta3 in the presence of botrocetin and ristocetin, respectively. J Thromb Haemost 4: 1367–1378.

Min, G., H. Wang, T. T. Sun and X. P. Kong (2006). Structural basis for tetraspanin functions as revealed by the cryo-EM structure of uroplakin complexes at 6-A resolution. J Cell Biol 173: 975–983.

Noy, P. J., J. Yang, J. S. Reyat, A. L. Matthews, A. E. Charlton, J. Furmston, D. A. Rogers, G. E. Rainger and M. G. Tomlinson (2016). TspanC8 Tetraspanins and A Disintegrin and Metalloprotease 10 (ADAM10) Interact via Their Extracellular Regions: EVIDENCE FOR DISTINCT BINDING MECHANISMS FOR DIFFERENT TspanC8 PROTEINS. J Biol Chem 291: 3145–3157. 10.1074/jbc.M115.703058.

Prox, J., M. Willenbrock, S. Weber, T. Lehmann, D. Schmidt-Arras, R. Schwanbeck, P. Saftig and M. Schwake (2012). Tetraspanin15 regulates cellular trafficking and activity of the ectodomain sheddase ADAM10. Cell Mol Life Sci 69: 2919–2932. 10.1007/s00018-012-0960-2.

Ran, F. A., P. D. Hsu, J. Wright, V. Agarwala, D. A. Scott and F. Zhang (2013). Genome engineering using the CRISPR-Cas9 system. Nat Protoc 8: 2281–2308. 10.1038/nprot.2013.143.

Reiss, K., T. Maretzky, A. Ludwig, T. Tousseyn, B. de Strooper, D. Hartmann and P. Saftig (2005). ADAM10 cleavage of N-cadherin and regulation of cell-cell adhesion and beta-catenin nuclear signalling. EMBO J 24: 742–752. 10.1038/sj.emboj.7600548.

Reyat, J. S., M. Chimen, P. J. Noy, J. Szyroka, G. E. Rainger and M. G. Tomlinson (2017). ADAM10-Interacting Tetraspanins Tspan5 and Tspan17 Regulate VE-Cadherin Expression and Promote T Lymphocyte Transmigration. J Immunol 199: 666–676. 10.4049/jimmunol.1600713.

Reyat, J. S., M. G. Tomlinson and P. J. Noy (2017). Utilizing Lentiviral Gene Transfer in Primary Endothelial Cells to Assess Lymphocyte-Endothelial Interactions. Methods Mol Biol 1591: 155–168. 10.1007/978-1-4939-6931-9_11.

Rubinstein, E., S. Charrin and M. G. Tomlinson (2013). Organisation of the tetraspanin web. Tetraspanins. F. Berditchevski and E. Rubinstein. Dordrecht, Springer: 47–90.

Sahin, U., G. Weskamp, K. Kelly, H. M. Zhou, S. Higashiyama, J. Peschon, D. Hartmann, P. Saftig and C. P. Blobel (2004). Distinct roles for ADAM10 and ADAM17 in ectodomain shedding of six EGFR ligands. Journal of Cell Biology 164: 769–779.

Saint-Pol, J., M. Billard, E. Dornier, E. Eschenbrenner, L. Danglot, C. Boucheix, S. Charrin and E. Rubinstein (2017). New insights into the tetraspanin Tspan5 using novel monoclonal antibodies. J Biol Chem 292: 9551–9566. 10.1074/jbc.M116.765669.

Saint-Pol, J., E. Eschenbrenner, E. Dornier, C. Boucheix, S. Charrin and E. Rubinstein (2017). Regulation of the trafficking and the function of the metalloprotease ADAM10 by tetraspanins. Biochem Soc Trans 45: 937–944. 10.1042/BST20160296.

Seipold, L., H. Altmeppen, T. Koudelka, A. Tholey, P. Kasparek, R. Sedlacek, M. Schweizer, J. Bar, M. Mikhaylova, M. Glatzel and P. Saftig (2018). In vivo regulation of the A disintegrin and metalloproteinase 10 (ADAM10) by the tetraspanin 15. Cell Mol Life Sci 75: 3251–3267. 10.1007/s00018-018-2791-2.

Shah, J., F. Rouaud, D. Guerrera, E. Vasileva, L. M. Popov, W. L. Kelley, E. Rubinstein, J. E. Carette, M. R. Amieva and S. Citi (2018). A Dock-and-Lock Mechanism Clusters ADAM10 at Cell-Cell Junctions to Promote alpha-Toxin Cytotoxicity. Cell Rep 25: 2132–2147 e2137. 10.1016/j.celrep.2018.10.088.

Sidahmed-Adrar, N., J. F. Ottavi, N. Benzoubir, T. Ait Saadi, M. Bou Saleh, P. Mauduit, C. Guettier, C. Desterke and F. Le Naour (2019). Tspan15 Is a New Stemness-Related Marker in Hepatocellular Carcinoma. Proteomics: e1900025. 10.1002/pmic.201900025.

Sievers, F., A. Wilm, D. Dineen, T. J. Gibson, K. Karplus, W. Li, R. Lopez, H. McWilliam, M. Remmert, J. Soding, J. D. Thompson and D. G. Higgins (2011). Fast, scalable generation of high-quality protein multiple sequence alignments using Clustal Omega. Mol Syst Biol 7: 539. 10.1038/msb.2011.75.

Tang, Z., B. Kang, C. Li, T. Chen and Z. Zhang (2019). GEPIA2: an enhanced web server for large-scale expression profiling and interactive analysis. Nucleic Acids Res 47: W556–W560. 10.1093/nar/gkz430.

Termini, C. M. and J. M. Gillette (2017). Tetraspanins Function as Regulators of Cellular Signaling. Front Cell Dev Biol 5: 34. 10.3389/fcell.2017.00034.

Thul, P. J., L. Akesson, M. Wiking, D. Mahdessian, A. Geladaki, H. Ait Blal, T. Alm, A. Asplund, L. Bjork, L. M. Breckels, A. Backstrom, F. Danielsson, L. Fagerberg, J. Fall, L. Gatto, C. Gnann, S. Hober, M. Hjelmare, F. Johansson, S. Lee, C. Lindskog, J. Mulder, C. M. Mulvey, P. Nilsson, P. Oksvold, J. Rockberg, R. Schutten, J. M. Schwenk, A. Sivertsson, E. Sjostedt, M. Skogs, C. Stadler, D. P. Sullivan, H. Tegel, C. Winsnes, C. Zhang, M. Zwahlen, A. Mardinoglu, F. Ponten, K. von Feilitzen, K. S. Lilley, M. Uhlen and E. Lundberg (2017). A subcellular map of the human proteome. Science 356. 10.1126/science.aal3321.

Tomlinson, M. G., S. D. Calaminus, O. Berlanga, J. M. Auger, T. Bori-Sanz, L. Meyaard and S. P. Watson (2007). Collagen promotes sustained glycoprotein VI signaling in platelets and cell lines. J Thromb Haemost 5: 2274–2283.

Tomlinson, M. G., T. Hanke, D. A. Hughes, A. N. Barclay, E. Scholl, T. Hünig and M. D. Wright (1995). Characterization of mouse CD53: epitope mapping, cellular distribution and induction by T cell receptor engagement during repertoire selection. Eur J Immunol 25: 2201–2206.

Tomlinson, M. G., A. F. Williams and M. D. Wright (1993). Epitope mapping of anti-rat CD53 monoclonal antibodies. Implications for the membrane orientation of the transmembrane 4 superfamily. Eur J Immunol 23: 136–140.

Tusher, V. G., R. Tibshirani and G. Chu (2001). Significance analysis of microarrays applied to the ionizing radiation response. Proc Natl Acad Sci U S A 98: 5116–5121. 10.1073/pnas.091062498.

Uhlen, M., C. Zhang, S. Lee, E. Sjostedt, L. Fagerberg, G. Bidkhori, R. Benfeitas, M. Arif, Z. Liu, F. Edfors, K. Sanli, K. von Feilitzen, P. Oksvold, E. Lundberg, S. Hober, P. Nilsson, J. Mattsson, J. M. Schwenk, H. Brunnstrom, B. Glimelius, T. Sjoblom, P. H. Edqvist, D. Djureinovic, P. Micke, C. Lindskog, A. Mardinoglu and F. Ponten (2017). A pathology atlas of the human cancer transcriptome. Science 357. 10.1126/science.aan2507.

van Deventer, S. J., V. E. Dunlock and A. B. van Spriel (2017). Molecular interactions shaping the tetraspanin web. Biochem Soc Trans 45: 741–750. 10.1042/BST20160284.

Virreira Winter, S., A. Zychlinsky and B. W. Bardoel (2016). Genome-wide CRISPR screen reveals novel host factors required for Staphylococcus aureus alpha-hemolysin-mediated toxicity. Sci Rep 6: 24242. 10.1038/srep24242.

Wetzel, S., L. Seipold and P. Saftig (2017). The metalloproteinase ADAM10: A useful therapeutic target? Biochim Biophys Acta. 10.1016/j.bbamcr.2017.06.005.

Wilson, E., K. Leszczynska, N. S. Poulter, F. Edelmann, V. A. Salisbury, P. J. Noy, A. Bacon, J. Z. Rappoport, J. K. Heath, R. Bicknell and V. L. Heath (2014). RhoJ interacts with the GIT-PIX complex and regulates focal adhesion disassembly. J Cell Sci 127: 3039–3051. 10.1242/jcs.140434.

Zhang, B., Z. Zhang, L. Li, Y. R. Qin, H. Liu, C. Jiang, T. T. Zeng, M. Q. Li, D. Xie, Y. Li, X. Y. Guan and Y. H. Zhu (2018). TSPAN15 interacts with BTRC to promote oesophageal squamous cell carcinoma metastasis via activating NF-kappaB signaling. Nat Commun 9: 1423. 10.1038/s41467-018-03716-9.

Zhou, J., T. Fujiwara, S. Ye, X. Li and H. Zhao (2014). Downregulation of Notch modulators, tetraspanin 5 and 10, inhibits osteoclastogenesis in vitro. Calcif Tissue Int 95: 209–217. 10.1007/s00223-014-9883-2.

Zimmerman, B., B. Kelly, B. J. McMillan, T. C. Seegar, R. O. Dror, A. C. Kruse and S. C. Blacklow (2016). Crystal Structure of a Full-Length Human Tetraspanin Reveals a Cholesterol-Binding Pocket. Cell 167: 1041–1051. 10.1016/j.cell.2016.09.056.

Zuidscherwoude, M., F. Gottfert, V. M. Dunlock, C. G. Figdor, G. van den Bogaart and A. B. van Spriel (2015). The tetraspanin web revisited by super-resolution microscopy. Sci Rep 5: 12201. 10.1038/srep12201.

